# CK2 inhibits TIMELESS nuclear export and modulates CLOCK transcriptional activity to regulate circadian rhythms

**DOI:** 10.1101/2020.06.12.148825

**Authors:** Yao D. Cai, Yongbo Xue, Cindy C. Truong, Jose Del Carmen-Li, Christopher Ochoa, Jens T. Vanselow, Katherine A. Murphy, Ying H. Li, Xianhui Liu, Ben L. Kunimoto, Haiyan Zheng, Caifeng Zhao, Yong Zhang, Andreas Schlosser, Joanna C. Chiu

**Author notes:** **Corresponding author**: Joanna C. Chiu,. **Author Contributions:** J.C.C., Y.D.C., A.S. designed research; Y.D.C., Y.X., C.C.T., J.D.C, C.O., K.A.M., Y.H.L., B.L.K., J.T.V., H.Z., C.Z. performed research and analyzed data; Y.D.C., J.C.C., Y.X., Y.Z., A.S. contributed to critical interpretation of the data; X.H.L., K.A.M. generated reagents; Y.D.C, A.S., and J.C.C. wrote the paper.

## Abstract

Circadian clocks orchestrate daily rhythms in organismal physiology and behavior to promote optimal performance and fitness. In *Drosophila*, key pacemaker proteins PERIOD (PER) and TIMELESS (TIM) are progressively phosphorylated to perform phase-specific functions. Whereas PER phosphorylation has been extensively studied, systematic analysis of site-specific TIM phosphorylation is lacking. Here, we identified phosphorylation sites of PER-bound TIM by mass spectrometry, given the importance of TIM as a modulator of PER function in the oscillator. Among the twelve TIM phosphorylation sites we identified, at least two of them are critical for circadian timekeeping as mutants expressing non-phosphorylatable mutations exhibit altered behavioral rhythms. In particular, we observed that CK2-dependent phosphorylation of TIM(S1404) promotes nuclear accumulation of PER-TIM heterodimers by inhibiting the interaction of TIM and nuclear export component, Exportin 1 (XPO1). We postulate that proper level of nuclear PER-TIM accumulation is necessary to facilitate kinase recruitment for the regulation of daily phosphorylation rhythm and phase-specific transcriptional activity of CLOCK (CLK). Our results highlight the contribution of phosphorylation-dependent nuclear export of PER-TIM heterodimers to the maintenance of circadian periodicity and identify a new mechanism by which the negative elements of the circadian oscillator (PER-TIM) regulate the positive elements (CLK-CYC). Finally, since the molecular phenotype of *tim*(S1404A) non-phosphorylatable mutant exhibits remarkable similarity to that of a mutation in human *timeless* that underlies Familial Advanced Sleep Phase Syndrome (FASPS), our results revealed an unexpected parallel between the functions of *Drosophila* and human TIM and may provide new insights into the molecular mechanisms underlying human FASPS.

**Significance Statement:** Phosphorylation is a central mechanism important for the regulation of circadian physiology across organisms. The molecular oscillator is composed of pacemaker proteins that undergo elaborate phosphorylation programs to regulate phase-specific functions. In *Drosophila*, phosphorylation of TIMELESS (TIM) has been recognized as critical for its function in the oscillator, but a systematic analysis of TIM phosphorylation is lacking. Here, we identified twelve *Drosophila* TIM phosphorylation sites by mass spectrometry and showed that phosphorylation at TIM(S1404) is necessary for maintaining 24-hour rhythms. Finally, since the molecular phenotype of *tim*(S1404A) non-phosphorylatable fly mutant exhibits remarkable similarity to that of a mutation in human *timeless* that underlies FASPS, our results may provide new insights into the molecular underpinnings of human FASPS.

## Introduction

Circadian rhythms have been observed in all domains of life and are driven by a network of cellular molecular clocks in animals (1–3). These molecular clocks are entrained by environmental time cues, such as light-dark (LD) cycles, to control daily rhythms in physiology and behavior. One conserved feature of molecular clocks within the animal kingdom is their reliance on key clock proteins that are organized in transcriptional translational feedback loops (TTFL) (3, 4). In *Drosophila*, these key pacemaker proteins are the positive elements, CLOCK (CLK) and CYCLE (CYC), and the negative elements, PERIOD (PER) and TIMELESS (TIM). During the day, CLK-CYC heterodimers activate the transcription of *per*, *tim* and other clock-controlled genes (ccgs) (5). The accumulation of PER and TIM proteins is delayed by a number of post-transcriptional and post-translational mechanisms (6–10) until early night when PER and TIM attain high enough levels to form heterodimeric complexes in the cytoplasm prior to nuclear entry (11, 12). Nuclear PER, likely still in complex with TIM, promotes the repression of the circadian transcriptome by inhibiting CLK-CYC transcriptional activity and removing them from clock genes before its degradation in the late night and early day (13–17).

Although TIM itself cannot repress CLK-CYC transcriptional activity, it is essential to the molecular clock because it maintains rhythmic PER protein expression (18–20). Constitutive cytoplasmic PER in *tim-null* mutants (21) as well as in *tim* mutants with defective nuclear entry (22, 23) abolishes rhythms in the molecular clock and consequently dampens behavioral rhythms. An early study suggested that TIM physically associates to the PER cytoplasmic localization domain (CLD) and thus blocks cytoplasmic retention (11). A subsequent study showed that TIM actively facilitates PER nuclear entry by acting as the primary cargo of Importin α1 (IMPα1)-dependent nuclear import mechanisms and cotransport PER into the nucleus (12). Finally, TIM is suggested to counteract the effect of DOUBLETIME (DBT) kinases in preventing PER nuclear entry (20). Taken together, TIM is proposed to promote the nuclear entry of PER.

Phosphorylation has also been implicated in regulating PER-TIM nuclear entry. The function of casein kinase 1α(CK1α), casein kinase 2 (CK2), SHAGGY (SGG) and DBT have been investigated in this context. DBT has been observed to phosphorylate PER and prevent nuclear translocation (20). This regulatory step has recently been shown to be antagonized by CK1α-dependent phosphorylation of either PER or DBT (24). The role of CK2 and SGG in regulating PER-TIM subcellular localization has received relatively more attention. Early studies suggest that SGG and CK2 phosphorylate both PER and TIM to promote nuclear translocation (25–27). Subsequent studies indicate that PER may be the primary target of CK2 and SGG to control subcellular localization (28, 29). However, more recent studies indicate that perhaps CK2 regulates nuclear entry of PER-TIM by phosphorylating TIM (30, 31). In addition to kinases, protein phosphatase 1 (PP1) and protein phosphatase 2A (PP2A) have also been shown to influence PER-TIM nuclear accumulation (32, 33).

In addition to its role in facilitating PER nuclear entry, TIM has been observed to mediate light resetting and circadian entrainment because of its light-induced degradation (34, 35). Upon light exposure, the blue light photoreceptor CRYPTOCHROME (CRY) undergoes a conformational change and binds to TIM (36–37). The E3 ubiquitin ligase JETLAG (JET) then collaborates with CRY to promote rapid proteasomal degradation of TIM (36, 38, 39). Phosphorylation of yet uncharacterized tyrosine residues has been proposed to be required for degradation (40). Finally, light-induced TIM degradation further promotes PER turnover, which function to reset and entrain the molecular clock (35).

Significant progress has been made in examining the function of site-specific PER phosphorylation, enabling in-depth mechanistic understanding of post-translational regulation of PER subcellular localization, repressor activity, and degradation to generate a 24-hr rhythm (28, 29, 41–45). On the other hand, the relative dearth of studies that characterize site-specific functions of TIM phosphorylation (22, 31, 46) remains a significant obstacle to fully understand the regulation of circadian rhythms via post-translational regulation of TIM and PER-TIM complexes. In this study, we used mass spectrometry proteomics to identify phosphorylation sites of PER-bound TIM proteins purified from *Drosophila* heads. We found that loss of phosphorylation at some of these TIM residues resulted in altered circadian behavioral rhythms. In particular, impaired CK2-dependent phosphorylation at TIM(S1404) resulted in a ~1.7-hr period-shortening phenotype. By analyzing the molecular clock of non-phosphorylatable *tim*(S1404A) and phosphomimetic *tim*(S1404D) mutants, we provide evidence supporting the importance of TIM(S1404) phosphorylation in promoting TIM nuclear retention by reducing its interaction with Exportin 1 (XPO1), an important component of the nuclear export machinery. Interestingly, decreased nuclear localization of TIM in *tim*(S1404A) mutant flies not only reduces the abundance of nuclear PER and TIM proteins, but also dampens the daily rhythms in CLK phosphorylation. We reasoned this is caused by changes in the abundance of kinases recruited by the PER-TIM complexes to phosphorylate CLK. Consequently, this leads to phase advance of CLK occupancy rhythms at circadian promoters, which manifests into shortening of molecular and behavioral rhythms. Based upon these findings, we propose a model describing the mechanism by which CK2-dependent TIM(S1404) phosphorylation regulates PER-TIM nuclear accumulation and CLK-CYC activity to regulate circadian rhythms.

## Results

### Mass Spectrometry analysis identifies TIM phosphorylation sites in PER-TIM heterodimers

Comprehensive mapping of TIM phosphorylated sites have not been performed despite previous studies indicating that phosphorylation is important for the phase-specific regulation of TIM function (25–27, 31, 40, 47). This hinders further understanding of the mechanisms by which site-specific TIM phosphorylation regulates the molecular clock. Since TIM interacts with PER to achieve its role in circadian timekeeping, we aimed to identify phosphorylated TIM residues in the PER-TIM heterodimeric complex. Previously, our group identified PER phosphorylation sites by purifying PER from fly heads using affinity purification followed by mass spectrometry (MS) analysis (48). We observed that TIM was copurified with PER (ZT1, 3, 12, 16, 20, 23.5) and took the opportunity to identify PER-bound TIM phosphorylation sites. The MS data from multiple time-points were pooled to identify TIM phosphorylation sites qualitatively. We expect that some of these phosphorylation events may be critical in regulating PER-TIM interactions and the function of PER-TIM heterodimer in the molecular clock.

We identified 12 TIM phosphorylation sites, some of which are located in previously characterized functional domains (Fig. 1A and Table S1). These include S568, which is in PER Binding Domain 1 (PER BD1) and the NLS (nuclear localization signal) (11), S891 in PER Binding Domain 2 (PER BD 2) (11), S1389, S1393, S1396, and S1404 in the Cytoplasmic Localization Domain (CLD) (11). Based on the location of these phosphorylation sites, we reasoned that they may regulate the subcellular localization and phase-specific functions of PER-TIM heterodimers. We also identified a number of other phosphorylated residues that are located in regions of TIM proteins without characterized functions. Finally, although not the central focus of this study, we determined that TIM, like PER and CLK (48–50), is O-GlcNAcylated at multiple residues (Table S1).

**Figure 1.**
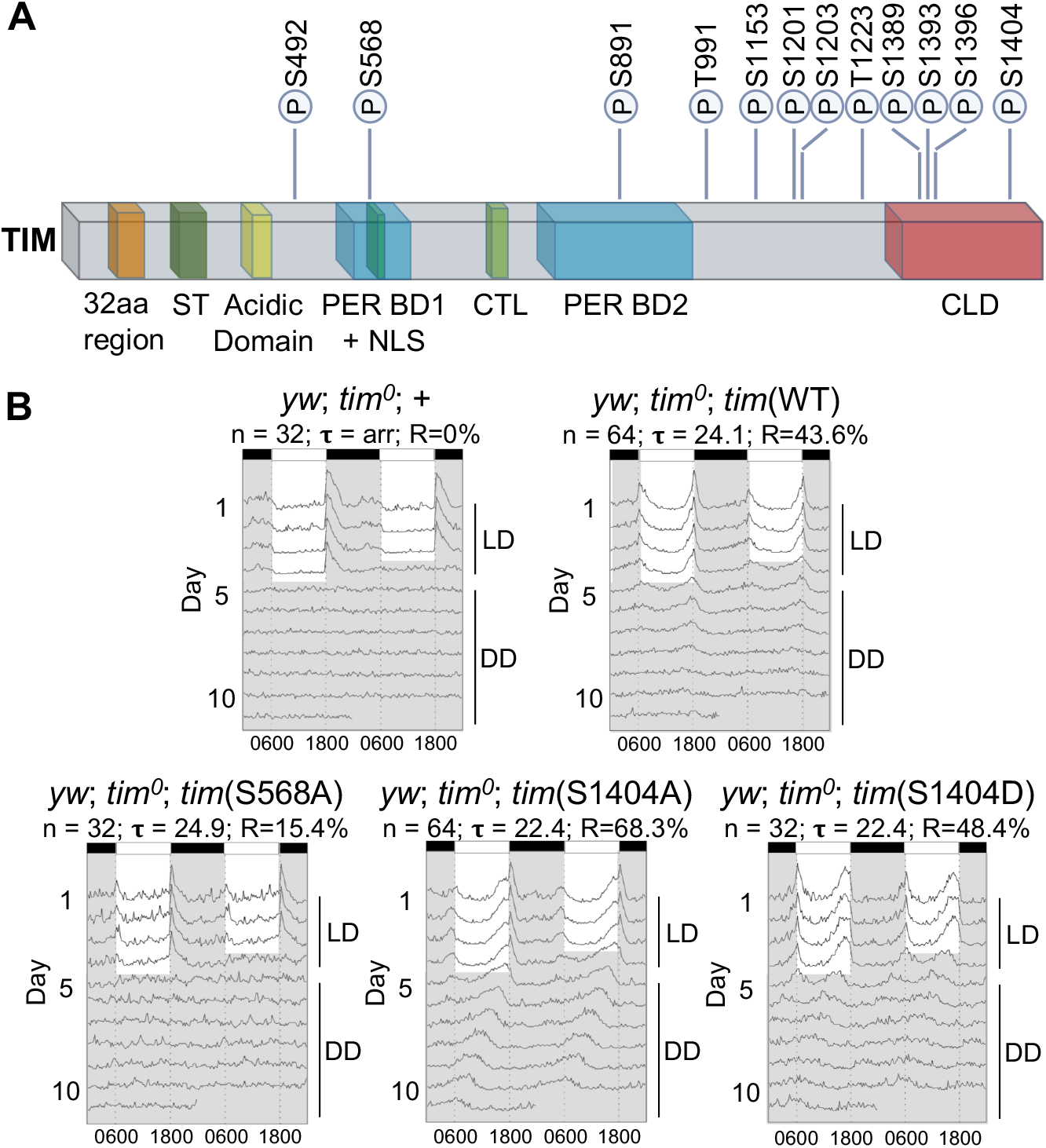
Daily locomotor activity rhythms are altered in TIM phosphorylation site mutants. (A) Schematic showing phosphorylation sites mapped onto TIM functional domains. All amino acid numbering is based on the L-TIM_1421_ isoform (81–83). Previously described domains of TIM: 32 amino acid region (aa 260-291) (81), also known as serine-rich domain (SRD, aa 260-292 (47); serine/threonine (ST)-rich region (aa 293-312) (31); acidic domain (aa 383-412) (81); PER binding domain 1 (PER BD1) (aa 536-610) (11); nuclear localization sequence (NLS) (aa 558-583) (11); C-terminal tail-like sequence (CTL) (aa640-649) (37); PER binding domain 2 (PER BD2) (aa 747-946) (11); cytoplasmic localization domain (CLD) (aa 1261-1421) (11). Corresponding PEAKS Studio scores of modified peptides are shown in Table S1. (B) Double-plotted actograms of *yw*; *tim^0^* flies carrying transgenes for site-specific TIM phosphorylation mutation generated using FaasX. n represents the sample size for behavioral assay. Tau (**τ**) represents the average period length of the indicated group of flies in DD. R represents percentage of flies that are rhythmic. Flies were entrained for 4 days in LD and then switched to 7 days of constant darkness, DD.

### Transgenic flies expressing non-phosphorylatable TIM variants display altered locomotor activity rhythms

To determine if the TIM phosphorylation sites we identified play important roles in circadian timekeeping, we generated transgenic fly lines each expressing one or a cluster of non-phosphorylatable S/T to A mutations. We prioritized our efforts by focusing on sites with high MS probability score and/or those located in characterized functional domains. *p*{*tim*(X)-3XFLAG-6XHIS} transgenes (X represents the S/T to A TIM mutation or WT TIM) were crossed into *tim^0^* genetic background (51) such that only transgenic *tim* was expressed. First, we evaluated daily locomotor activity rhythms of *tim* transgenic flies as activity rhythm is a reliable behavioral output of the *Drosophila* circadian clock (52). Flies were entrained for 3 days in 12 hours light and 12 hours darkness cycles (herein referred as LD cycles) followed by 7 days in constant darkness (DD) to monitor free-running rhythm. We observed that homozygous *tim*(WT) flies exhibited behavioral rhythm with a ~24-hr period, indicating that the arrhythmic *tim^0^* mutation was rescued by the *tim*(WT) transgene (Fig. 1B, S1A, and Table S2). Although the behavioral rhythmicity of *tim*(WT) is relatively low (43.6%) compared to what is normally observed in wild type flies or *per^0^* rescues (e.g. 48), the rhythmicity is notably higher than rescue of *tim^0^* flies by driving expression of *tim* cDNA using *tim-*Gal4 (22, 52), and comparable to the extent of *tim^0^* rescue observed as previously described (31). Among the transgenic lines expressing TIM variants, *tim*(S1404A) was the only genotype that exhibited a clear period-shortening phenotype (~1.7 hr shorter), while *tim*(S568A) flies were notably more arrhythmic when compared to *tim*(WT).

Among the two phosphorylation sites that resulted in notable changes in behavioral rhythms when mutated, we decided to proceed first with the functional characterization of TIM(S1404) as it is (i) highly conserved in animals (Fig. S2A and S2B), (ii) predicted to be phosphorylated by CK2, a known clock kinase, (iii) also phosphorylated in the monarch butterfly TIM protein as determined by MS analysis (Fig. S2C), and (iv) located within TIM CLD (Fig. 1A). For this reason, we generated an additional transgenic fly line expressing a phosphomimetic S1404D TIM variant to complement the analysis of *tim*(S1404A) mutants. Interestingly, *tim*(S1404D) flies also exhibited shortened period, similar to what was observed in *tim*(S1404A) flies (Fig. 1B and Table S2). Although it is logical to assume that non-phosphorylatable and phosphomimetic mutations should result in opposite phenotypes (e.g. 31, 41), that is often not the case as observed in previous phosphorylation studies (e.g. 28, 42, 45). Furthermore, although both *tim*(S1404A) and *tim*(S1404D) mutants exhibit period-shortening at the behavioral level, the underlying molecular mechanisms that underlie their phenotypes may differ (e.g. 45).

### TIM(S1404) phosphorylation promotes TIM nuclear accumulation

Since TIM S1404 residue is located in the CLD, we reasoned that S1404 phosphorylation may regulate TIM nuclear accumulation. To test our hypothesis, we monitored subcellular localization of TIM in *tim*(WT), *tim*(S1404A), and *tim*(S1404D) adult brain clock neurons from early to late night using whole-mount immunocytochemistry. These experiments were performed using flies entrained in LD cycles to preclude phase differences between the genotypes that are caused by alterations in period length. Costaining of TIM with pigment-dispersing factor (PDF) enabled the identification of PDF+ clock neurons (sLNvs and lLNvs) and demarcation of nuclear vs cytoplasmic compartments as PDF is expressed in the cytoplasm (53). In agreement with previous studies (53), we observed that the majority of TIM was cytoplasmic at ZT16 in *tim*(WT) flies but become progressively more nuclear from early to late night (Fig. 2A and 2B). In contrast, *tim*(S1404A) flies displayed significantly lower percentage of nuclear TIM (% nuclear TIM/total TIM) at late night (ZT20 and ZT22) (Fig. 2B), while phospho-mimetic *tim*(S1404D) mutants exhibited higher percentage of nuclear TIM over *tim*(WT) flies at ZT16 and ZT22.

**Figure 2.**
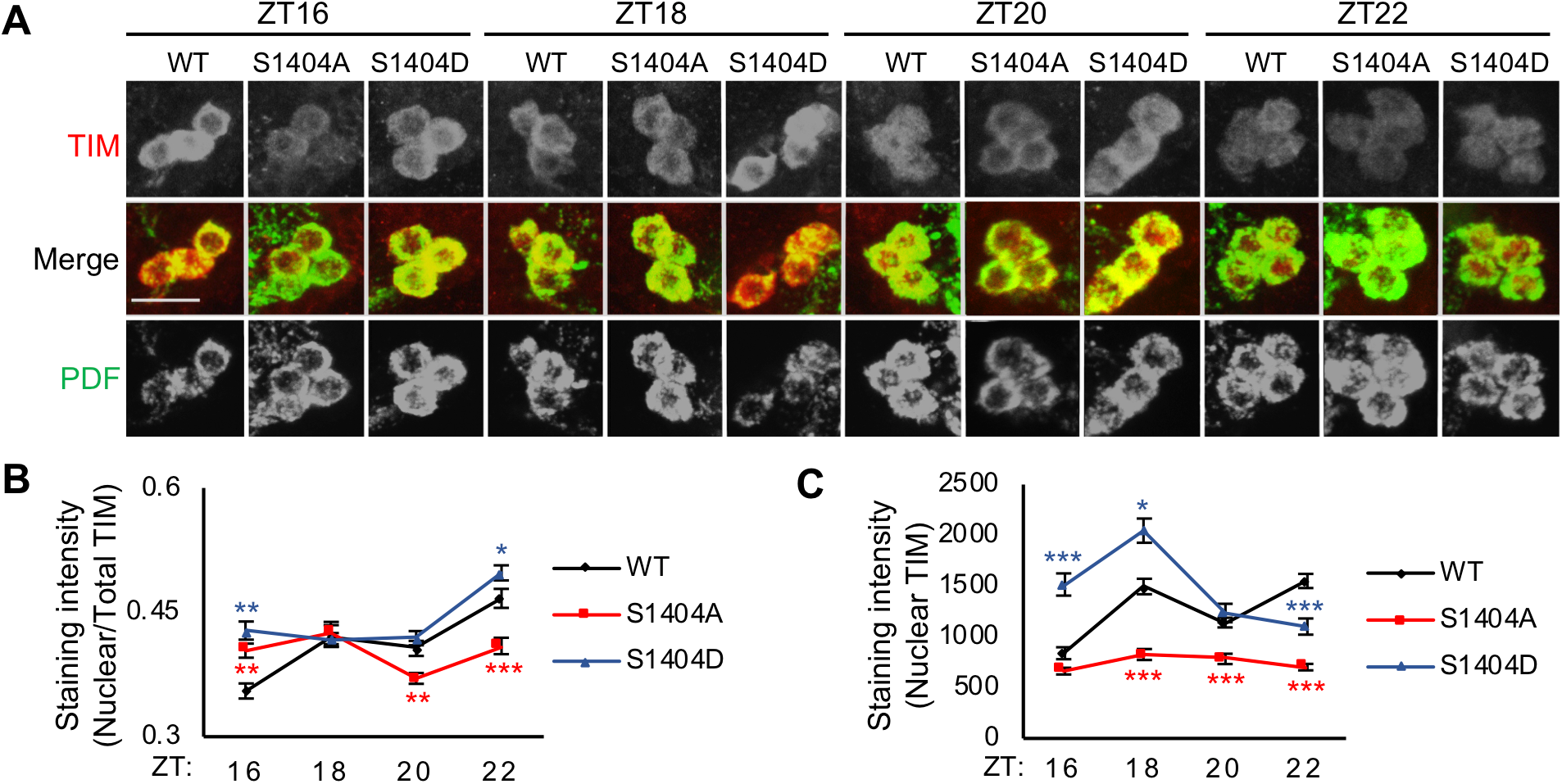
TIM nuclear accumulation is altered in *tim*(S1404A) and *tim*(S1404D) mutants. (A) Representative confocal images of sLNvs clock neurons in adult fly brains stained with α-TIM (red) and α-PDF (green). Single channels are shown in grey scale. Scale bar (merged image in WT ZT16) represents 10μm. Flies were entrained for 4 days in LD cycles and collected at the indicated times on LD4 for fixation and immunofluorescence analysis. (B) Line graph showing the fraction of nuclear TIM presented as nuclear TIM divided by total TIM in sLNvs. (C) Line graph showing nuclear TIM staining intensity in sLNvs. Error bars indicate SEM (n>30), ***p<0.001, **p<0.01, *p<0.05, Kruskal-Wallis test.

In addition to assessing the distribution of TIM in the nucleus vs. cytoplasm, we also monitored overall nuclear TIM abundance. We observed a substantially lower abundance of nuclear TIM in *tim*(S1404A) flies between ZT18 to ZT22, while that in *tim*(S1404D) flies appeared higher than *tim*(WT) at ZT16 to ZT18 but lower at ZT22 (Fig. 2A and 2C). Taken together, our results suggest that TIM(S1404) phosphorylation promotes TIM nuclear accumulation and the molecular mechanisms underlying the short-period behavioral phenotypes of *tim*(S1404A) and *tim*(S1404D) flies are likely different.

### TIM(S1404) phosphorylation regulates TIM-XPO1 interaction

A number of studies have established that PER-TIM nuclear accumulation is regulated by phosphorylation (20, 24, 28–31, 54). With the characterization of TIM nuclear import pathway (12) and functional NLS (23), phosphorylation has been thought to influence nuclear entry. However, the evidence that nuclear PER-TIM can be translocated back to the cytoplasm (11, 19) raise the possibility that phosphorylation may also regulate nuclear export to influence overall levels of nuclear accumulation. To determine whether S1404 phosphorylation regulates TIM nuclear entry or export, we first searched for potential NLS and nuclear export signal (NES) in the sequences adjacent to S1404 based on classical NLS/NES motifs (55, 56). Whereas we did not locate any sequences that resemble an NLS near S1404, we identified one putative NES (L1394-V1403) immediately adjacent to the S1404 residue (Fig. 3A). Previous studies suggest that phosphorylated residue(s) within or in close proximity to an NES can regulate protein nuclear-cytoplasmic distribution by modulating the binding of cargo protein and Chromosome Maintenance 1 (CRM1) (57, 58). CRM1 is the major export protein in mammals that facilitates the transport of proteins from the nucleus to the cytoplasm.

**Figure 3.**
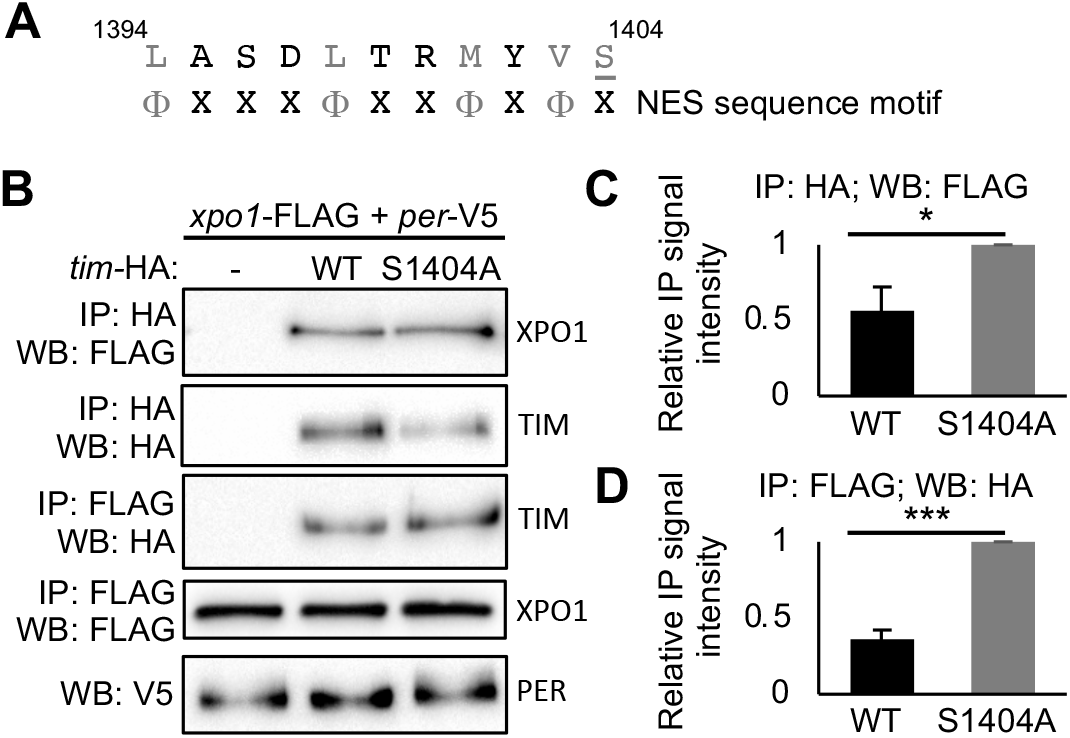
TIM(S1404) phosphorylation promotes TIM nuclear retention by compromising TIM-XPO1 interaction. (A) S1404 is located next to a putative TIM^NES^: L1394-V1403. S1404 is underlined and shown in grey. Classical NES sequence motif is previously investigated (57). Φ is hydrophobic amino acid (in grey): Leu, Val, Ile, Phe or Met; X is any amino acid. (B) Western blots showing reciprocal coimmunoprecipitations (coIPs) to examine the interactions of TIM(WT) or TIM(S1404A) to XPO1 in *Drosophila* S2 cells expressing pAc-*xpo1*-3XFLAG-6XHIS and pAc-*per*-V5 in the presence or absence of pAc-HA plasmids expressing *tim* variants. Protein extracts were directly analyzed by immunoblotting (α-V5 for PER) or immunoprecipitated with α-HA or α-FLAG resins to detect baits and interactors. (C and D) Bar graphs displaying quantification of reciprocal coIPs. Values for binding are normalized to amount of bait detected in the IPs and expressed as relative signal intensity (high value = 1). Error bars indicate ± SEM (n=4), ***p<0.001, *p<0.05, two-tailed Student’s t test.

We therefore tested whether TIM(S1404) phosphorylation reduces the interaction between TIM and XPO1, the *Drosophila* homolog of mammalian CRM1 by performing coimmunoprecipitation (coIP) assays using *Drosophila* S2 cells coexpressing *tim*(WT)-HA or *tim*(S1404A)-HA with *xpo1*-FLAG and *per*-V5. We observed significantly higher TIM(S1404A)-XPO1 interaction as compared to TIM(WT) when we pulled down TIM-HA and detected the presence of interacting XPO1 (Fig. 3B and 3C). We also performed the reciprocal coimmunoprecipitation, which yielded the same conclusion (Fig. 3B and 3D). Furthermore, we assayed the binding of TIM(S1404D) to XPO1 and observed that it was significantly lower when compared to that of TIM(S1404A) but similar to the levels for TIM(WT) (Fig. S3). The similarity between the interactions between TIM(WT)-XPO1 and TIM(S1404D)-XPO1 can be explained by the confirmation that TIM(S1404) is phosphorylated in TIM(WT) expressed in *Drosophila* S2 cells using a pS1404 phospho-specific antibody we generated for this study (Fig. S4A and S4B). Taken together, our results suggest that TIM(S1404) phosphorylation inhibits the nuclear export of TIM via the XPO1-dependent pathway.

### TIM(S1404) phosphorylation increases PER nuclear accumulation

Since TIM is necessary for promoting the nuclear entry of PER-TIM heterodimers (12, 22, 23), we next sought to determine whether PER nuclear accumulation is also altered in *tim*(S1404A) and *tim*(S1404D) mutants. We monitored subcellular localization of PER in adult clock neurons using the same method as described for TIM. As expected, the percent of nuclear PER (% nuclear PER/total PER) gradually increased from ZT16 to ZT22 in *tim*(WT) flies (Fig. 4A and 4B). In comparison, the percent of PER in the nucleus in *tim*(S1404A) mutants was significantly lower at ZT22 while that for *tim*(S1404D) mutant was significantly higher at ZT22. Furthermore, the overall abundance of nuclear PER was also significantly lower in *tim*(S1404A) mutants at ZT22 (Fig. 4A and 4C). Our results therefore support that alterations in TIM subcellular localization due to phosphorylation defect at TIM(S1404) impact subcellular localization of its heterodimeric partner PER.

**Figure 4.**
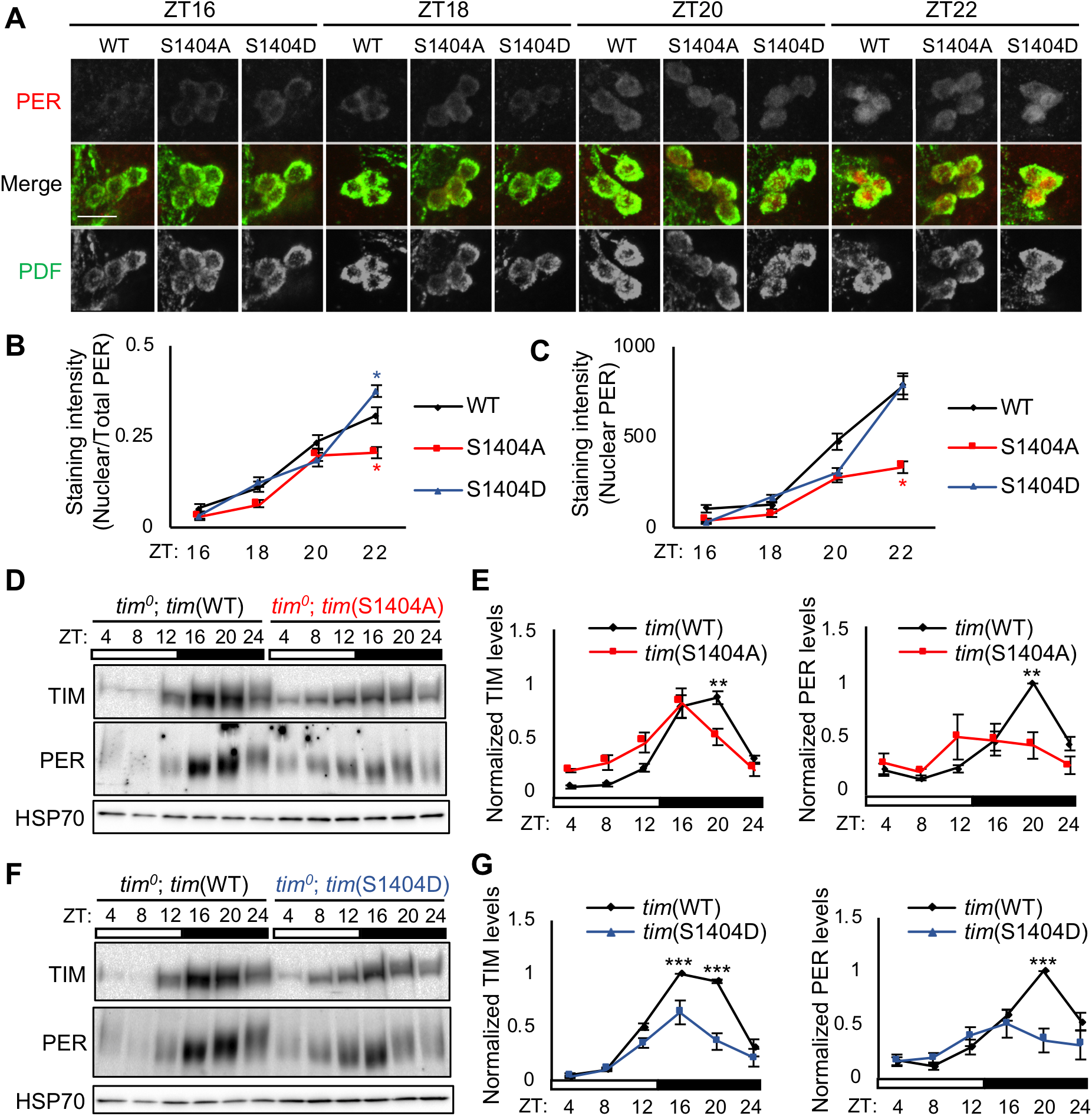
Altered TIM(S1404) phosphorylation influences PER nuclear accumulation. (A) Representative confocal images of sLNv clock neurons in adult fly brains stained with α-PER (red) and α-PDF (green). Single channels are shown in grey scale. Scale bar (merged image in WT ZT16) represents 10μm. Flies were entrained as described in Fig. 2A. (B) Line graph showing the fraction of nuclear PER in sLNvs presented as nuclear PER divided by total PER. (C) Line graph showing nuclear PER staining intensity in sLNvs. Error bars indicate ± SEM (n>27), *p<0.05, Kruskal-Wallis test. (D and F) Western blots comparing TIM and PER profiles in heads of (D) *tim*(WT) and *tim*(S1404A) flies or (F) *tim*(WT) and *tim*(S1404D) flies. Flies were entrained in LD cycles and collected on LD3 at indicated time-points (ZT). ⍺-HSP70 was used to indicate equal loading and for normalization. (E and G) Quantification of TIM and PER in (D) and (F). Error bars indicate ± SEM (n=3), ***p<0.001, **p<0.01, two-way ANOVA.

We next compared PER and TIM protein profiles in head extracts of WT and mutants to determine if altered nuclear accumulation affects their daily rhythms in protein abundance and phosphorylation state. Consistent with previous studies, newly synthesized PER and TIM in *tim*(WT) flies were hypophosphorylated between ZT 8 to ZT12 and became progressively more phosphorylated from early night to the following morning (24, 25, 59) (Fig. 4D and 4F). Daily rhythms in PER and TIM protein abundance were altered in *tim*(S1404A) mutants, as determined by Detection of Differential Rhythmicity (DODR) analysis (PER: p<0.05; TIM: p<0.01) (60). In congruence with the short period phenotype of *tim*(S1404A) flies, the peak phases of both PER and TIM rhythms advanced from ZT20 in *tim*(WT) flies to ZT16 (Fig. 4D and 4E) as calculated by Rhythmicity Analysis Incorporating Nonparametric methods (RAIN) (61). In addition, daily PER protein rhythmicity was dampened in *tim*(S1404A) mutants (WT: p<0.0001; S1404A: p= 0.1361, RAIN). This is likely caused by compromised nuclear accumulation of the PER-TIM proteins, which is clearly affecting their phosphorylation programs and is expected to impact their phase-specific functions. Furthermore, phase advance of *per* and *tim* mRNA, discussed in the next section, is also expected to contribute to changes in PER and TIM proteins rhythms.

In the case of the short period *tim*(S1404D) mutant, daily rhythms in PER and TIM were significantly altered as compared to *tim*(WT) (PER: p<0.05; TIM: p<0.05, DODR). The peak phase of PER advanced from ZT20 to ZT16 while that for TIM remained unchanged, as determined by RAIN (Fig. 4F and 5G). Moreover, we observed significant dampening of the daily rhythmicity of PER proteins in *tim*(S1404D) flies (WT: p<0.0001; S1404D: p=0.0720, RAIN). Daily rhythmicity of TIM in *tim*(S1404D) mutants was slightly dampened as compared to *tim*(WT) although still rhythmic (WT: p<0.0001; S1404D: p<0.0001, RAIN). Notably, the accelerated PER and TIM turnover in *tim*(S1404D) flies at night (Fig. 4G) is consistent with their increased nuclear localization at ZT22 (Fig. 2B and 4B). Together, our data suggests that TIM(S1404) phosphorylation promotes PER nuclear accumulation indirectly by increasing TIM nuclear retention.

### TIM(S1404) phosphorylation influences rhythmic CLK phosphorylation and occupancy at circadian promoters

We next examined whether reduced PER-TIM nuclear accumulation in *tim*(S1404A) mutants affects the output of the molecular oscillator by assaying the cycling of *per* and *tim* mRNAs. The daily rhythms in *per* and *tim* mRNAs in *tim*(WT) and *tim*(S1404A) mutants were significantly different (PER: p<0.001; TIM: p<0.001, DODR) (Fig. 5A). Given the reduction of nuclear PER to repress CLK-CYC transcriptional activity in *tim*(S1404A) flies at night, we expect *per* and *tim* mRNA levels in *tim*(S1404A) to be higher when compared to *tim*(WT) flies during the circadian repression phase (~ZT16-24) (16). Surprisingly, *per* and *tim* mRNA levels were substantially lower at ZT16 and ZT20 in the *tim*(S1404A) mutant. In addition, we observed a significant phase advance in *per* mRNAs in *tim*(S1404A) flies, in congruence with the short period phenotype of this mutant (WT: peak=ZT16; S1404A: peak=ZT12, RAIN). This resulted in significantly higher levels of *per* and *tim* mRNAs at ZT8 (Fig. 5A).

**Figure 5.**
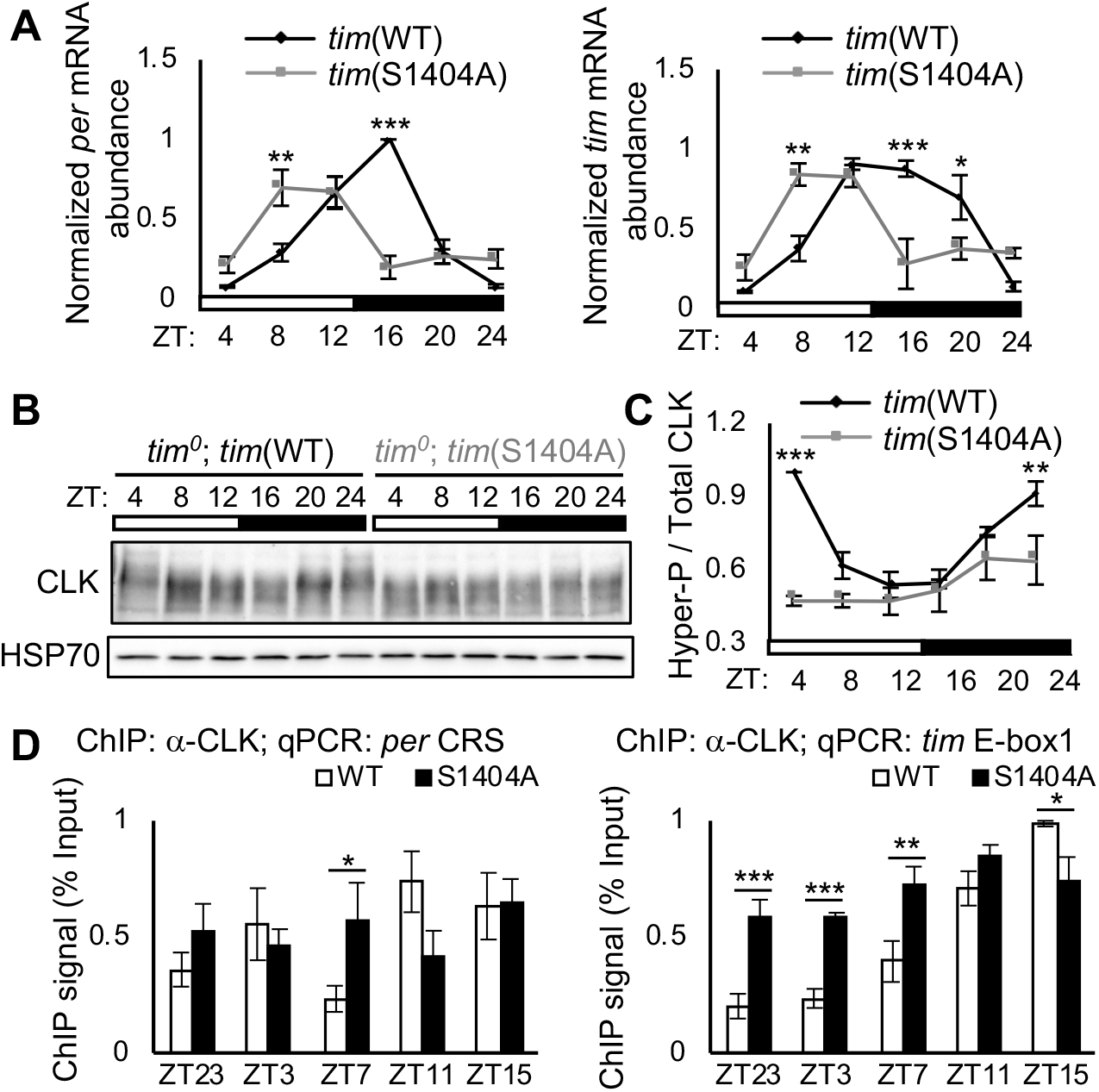
Reduced TIM nuclear retention in *tim*(S1404A) mutant leads to dampening of CLK phosphorylation rhythm and phase advance in CLK-activated transcriptional activation. (A) Steady state mRNA expression of *per* and *tim* in heads of *tim*(WT) and *tim*(S1404A) flies. Flies were entrained in LD cycles and collected on LD3 at indicated time-points (ZT) (n=3). (B) Western blots comparing CLK protein profiles in heads of *tim*(WT) and *tim*(S1404A) entrained and collected as in (A). ⍺-HSP70 was used to indicate equal loading and for normalization. (C) Quantification of hyperphosphorylated/total CLK. The top half of the CLK signal shown at ZT24 in *tim*(WT) flies (lane 6) is used as a reference to classify CLK isoforms as hyperphosphorylated (n=3). (D) ChIP assays using fly head extracts comparing CLK occupancy at *per* and *tim* promoters in *tim*(WT) and *tim*(S1404A) flies. CLK-ChIP signals were normalized to % input. ChIP signals for two intergenic regions were used for non-specific background deduction (n=4). Error bars indicate ± SEM, ***p<0.001, **p< 0.01, *p< 0.05, two-way ANOVA.

Our analysis of clock gene expression clearly suggests that the short period phenotype of *tim*(S1404A) is driven primarily by the phase advance of CLK transcriptional activity. But how does reduced nuclear accumulation of PER-TIM heterodimers leads to premature activation of CLK transcriptional activity? CLK transcriptional activity has previously been shown to correlate with CLK phosphorylation status (15, 62–67). Subsequent to nuclear translocation of PER-TIM heterodimers, kinases recruited by the PER-TIM complex have been proposed to phosphorylate CLK and inactivate its transcriptional activity (15, 62). Dephosphorylation by phosphatase then produce hypophosphorylated, transcriptionally active CLK in the following morning (68). Since TIM(S1404A) mutation reduces PER-TIM nuclear accumulation (Fig. 2B and 4B), we decided to examine its impact on the daily oscillation of CLK phosphorylation (Fig. 5B and 5C). We observed significant alteration in the daily rhythm of CLK phosphorylation in *tim*(S1404A) as compared to *tim*(WT) flies (p<0.001, DODR), despite no significant change in CLK abundance (Fig. S5A). Specifically, *tim*(S1404A) exhibited significant dampening in the daily CLK phosphorylation rhythms as compared to *tim*(WT) flies (WT: p<0.0001; S1404A: p= 0.1079, RAIN). In particular, significantly less hyperphosphorylated CLK isoforms were detected at ZT24/0 and ZT4 in *tim*(S1404A) flies, the time when CLK is predominantly hyperphosphorylated in *tim*(WT) flies (Fig. 5B and 5C). Since hypophosphorylated or intermediately phosphorylated CLK proteins have higher transcriptional activity (15, 63, 66), our results could explain the phase advance in CLK activation of *per* and *tim* expression in *tim*(S1404A) flies (Fig. 5A).

We then asked whether reduction in early morning CLK phosphorylation in *tim*(S1404A) mutants influences CLK occupancy at clock gene promoters and contributes to premature initiation of *per* and *tim* expression. We performed CLK chromatin immunoprecipitation (CLK-ChIP) followed by qPCR using extracts from adult fly heads (Fig. 5D). We observed significantly higher CLK occupancy at ZT7 at *per* CRS and multiple morning time-points at *tim* E-box in the *tim*(S1404A) mutant as compared to that in *tim*(WT) flies. Together, our data suggest that TIM(S1404) phosphorylation can impact daily rhythms in CLK phosphorylation status and transcriptional activity to regulate circadian timekeeping.

In the case of *tim*(S1404D) flies, although daily rhythms in *per* and *tim* mRNAs were not significantly different from *tim*(WT) flies (*per*: p=0.1742; *tim*: 0.4254, DODR), the peak phase of *per* mRNA was advanced from ZT16 to ZT12 (RAIN), and the repression of both *per* and *tim* appeared to occur earlier (Fig. S6A). This may contribute to the advanced peak phase in PER protein rhythms in *tim*(S1404D) flies (Fig. 4F and 4G). Since CLK protein abundance was not significantly altered in *tim*(S1404D) mutants (Fig. S5B), the advanced peak phase of *per* mRNA is likely a consequence of alteration in CLK phosphorylation rhythm. In agreement with our hypothesis, we observed a phase advance in the peak of hyperphosphorylated/total CLK in *tim*(S1404D) from ZT4 to ZT0/24 (RAIN), even though daily rhythms in CLK phosphorylation was not significantly altered (p=0.3140, DODR) (Fig. S6B and S6C).

### CK2 kinase phosphorylates TIM(S1404)

We next sought to identify the kinase that phosphorylates TIM(S1404). Based on KinasePhos 2.0 (69), CK2 is predicted with the highest probability to phosphorylate S1404 (Fig. 6A). To confirm this *in silica* prediction, we assayed TIM(S1404) phosphorylation in protein extracts of *Drosophila* S2 cells coexpressing *tim*-HA (WT or S1404A) with either the catalytic subunit of *ck2* (*ck2α*) or a dominant negative variant *ck2α*(*tik*) (26, 30). As expected, immunoblotting showed that TIM(pS1404) was significantly reduced when *tim*(WT)-HA was coexpressed with *ck2α*(*tik*) as compared to coexpression with *ck2α*(WT) (Fig. 6B, lanes 2-3, and 6C). Moreover, there was little to no α-pS1404 signal detected in *tim*(S1404A) (Fig. 6B, lanes 4-6, 6C), suggesting the α-pS1404 antibody is phosphospecific. To further validate the specificity of α-pS1404, we confirmed the reduction of α-pS1404 isoforms when TIM was immunoprecipitated and phosphatase-treated prior to immunoblotting (Fig. S4A, lanes 1-2, and S4B).

**Figure 6.**
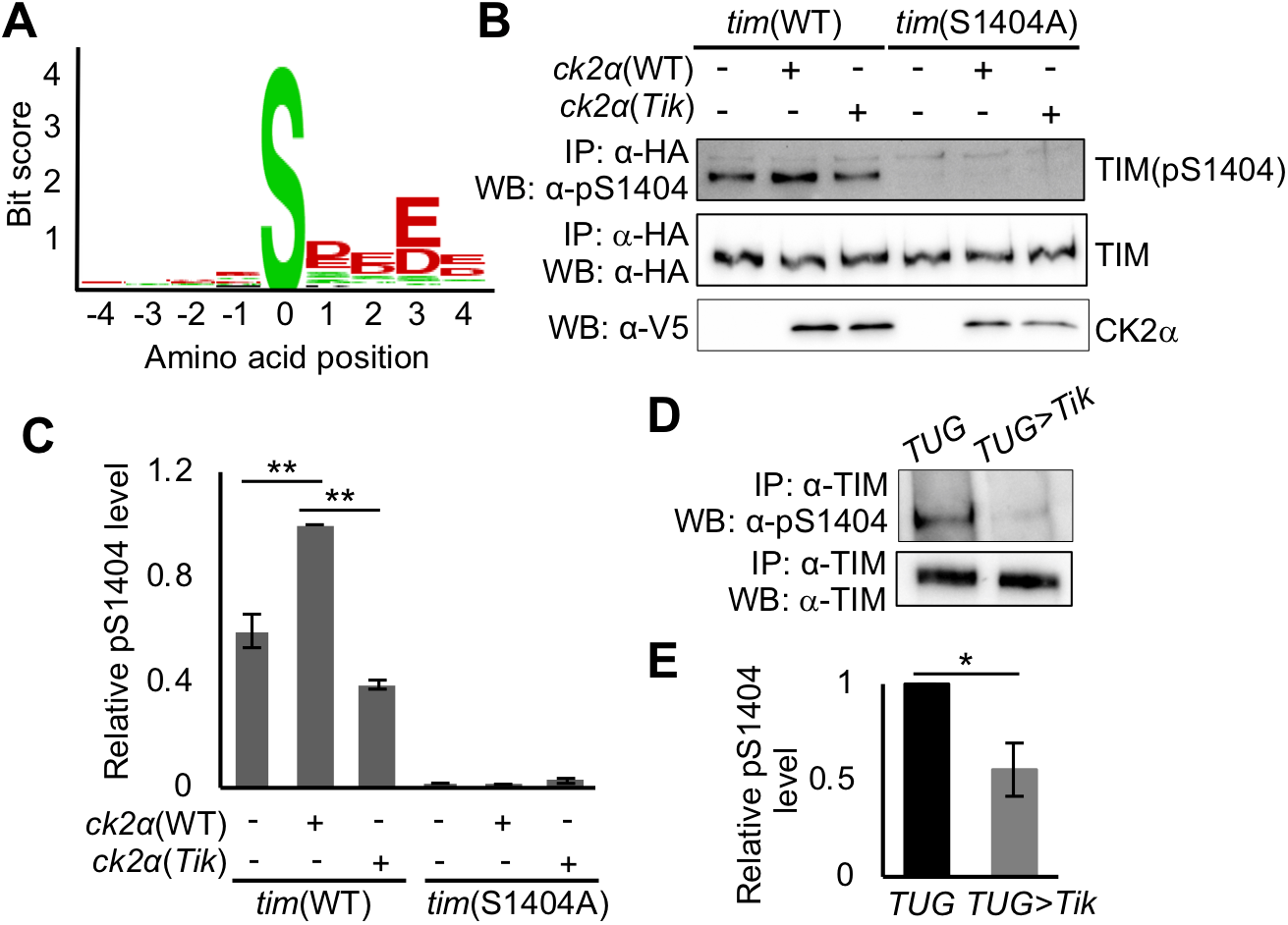
CK2 phosphorylates TIM(S1404). (A) CK2 consensus motifs generated by KinasePhos 2.0. S1404 corresponds to phosphoserine at amino acid position 0 (SVM Score=0.9581). (B) *Drosophila* S2 cells were transfected with pAc-*tim*(WT)-HA or pAc-*tim*(S1404A)-HA and co-transfected with an empty plasmid (pMT-V5-His), pMT-*ck2α*-V5, or pMT-*ck2α*(M161K E165D)-V5, referred to as *ck2α*(*tik*). Protein extracts were incubated with α-HA resin. Total TIM isoforms, TIM(pS1404), and CK2α protein levels were analyzed by Western Blotting with indicated antibodies. (C) Bar graph showing relative TIM pS1404 levels in (B) normalized to total TIM isoforms. Error bars indicate ± SEM (n=2), **p<0.01, two-way ANOVA. (D) Reduced pS1404 in flies overexpressing *ck2α*(*tik)* in *tim*-expressing cells (*TUG>tik*) as compared to parental control (*TUG*). Flies were entrained and collected on LD3 at ZT20. Fly head extracts were immunoprecipitated with α-TIM. TIM(pS1404) and total TIM isoforms were analyzed by Western blotting. (E) Bar graph showing relative pS1404 levels in (D), normalized to total TIM isoforms. Error bars indicate ± SEM (n=4), *p < 0.05, two-tailed Student’s t test.

We proceeded to test whether downregulating CK2 activity in flies reduces TIM(S1404) phosphorylation. First, we evaluated TIM(S1404) phosphorylation over a daily cycle and observed that phosphorylation at TIM(S1404) was detected at ZT16 and ZT20 in *tim*(WT) but absent in *tim*(S1404A) flies (Fig. S4C). Next, we genetically knocked down CK2 activity by overexpressing *ck2α*(*tik*) in clock neurons using the *tim*-UAS-Gal4 driver (*TUG*) (70). Head extracts from *TUG*>UAS-*ck2α*(*tik*) flies and parental controls collected at ZT20 were probed for S1404 phosphorylation. In concordance with the results in S2 cells, we observed a significant reduction in S1404 phosphorylation in *ck2α*(*tik*) overexpressing flies (Fig. 6D and 6E). Taken together, our results suggest that CK2 phosphorylates TIM(S1404) in *tim*-expressing clock neurons.

Finally, we performed immunocytochemistry in adult brain clock neurons to investigate the subcellular localization of CK2-dependent phosphorylation of TIM(pS1404). Based on western blotting results showing TIM(pS1404) signal at ZT16 in whole head extracts (Fig. S4C), abundant cytoplasmic localization of CK2 (26), and previous studies indicating the role of CK2 in promoting PER-TIM nuclear import (26, 27, 31, 47), we hypothesize that CK2 first phosphorylates TIM at S1404 and residues important for nuclear import in the cytoplasm, even though our results indicate that the function of TIM(S1404) phosphorylation is to inhibit nuclear export. In agreement with our observation from whole head extracts, we observed prominent TIM(pS1404) signal in clock neurons at ZT16 in *tim*(WT) but not in *tim*(S1404A) mutants (Fig. S4D).

## Discussion

To better understand the role of phosphorylation in regulating the function of the PER-TIM heterodimer, we identified multiple phosphorylation sites on PER-bound TIM proteins extracted from *Drosophila* tissues. After an initial behavioral screen of *tim* non-phosphorylatable mutants, we proceeded to characterize the function of TIM(S1404), which is located in the TIM CLD and is predicted to be phosphorylated by CK2, a known clock kinase. Leveraging the results from a series of molecular and behavioral analyses on transgenic flies expressing *tim*(S1404A) and *tim*(S1404D) mutants, we formulated a model describing the function of CK2-dependent TIM(S1404) phosphorylation in the molecular clock (Fig. 7). In wild type flies, TIM(S1404) is first phosphorylated by CK2 in the cytoplasm in early night. Our data does not rule out the possibility that CK2-dependent phosphorylation of TIM(S1404) continues after the entry of PER-TIM heterodimer into the nucleus around midnight. After nuclear entry, TIM(S1404) phosphorylation inhibits the interaction of TIM and the nuclear export machinery, thereby promoting nuclear accumulation of PER-TIM heterodimers. This facilitates timely CLK phosphorylation by kinases recruited by the PER-TIM complex to enhance circadian repression. The identity of kinase(s) that serve this role will need to be resolved in future investigations. Hyperphosphorylated CLK is then dephosphorylated by the CKA-PP2A complex (68) in the following morning to activate the next round of clock gene expression.

**Figure 7.**
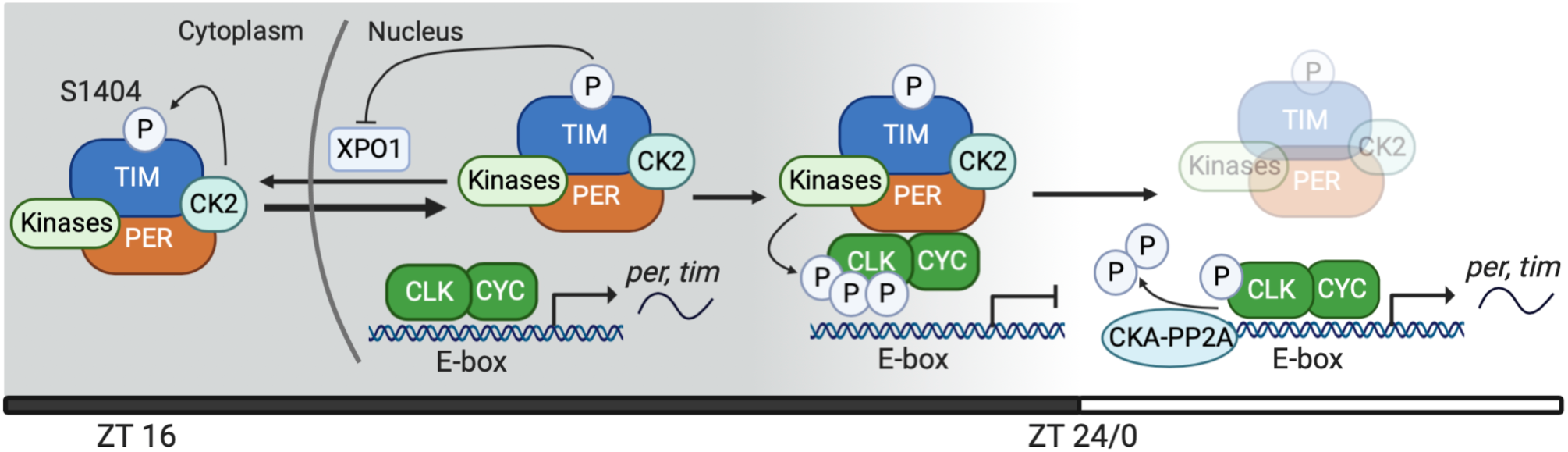
A model describing the function of TIM(S1404) phosphorylation in regulating the molecular clock. TIM(S1404) is phosphorylated by CK2 in the cytoplasm in early night. Upon entering the nucleus, phosphorylation at S1404 inhibits interaction of TIM and XPO1 and nuclear export of PER-TIM heterodimers, thereby promoting their nuclear accumulation. This allows kinase(s) bound to PER-TIM complex to phosphorylate CLK and remove CLK from circadian promoters. The kinase(s) responsible for this step is currently unknown. CKA-PP2A then dephosphorylates CLK and promotes the onset of CLK transcriptional activity in the next cycle (68). Other phosphorylation events on PER-TIM are not depicted for simplicity.

Our model is consistent with previous studies showing that PER-TIM-DBT complexes recruit as yet unknown kinases to phosphorylate CLK (67). It is also consistent with studies proposing that CK2 regulates PER function by phosphorylating TIM (31, 47). Since the S1404A mutation results in a short period phenotype, which is opposite to the period-lengthening effect of *ck2α^tik^* (26), our results highlight the complex functions of CK2 in regulating the molecular clock.

In *tim*(S1404A) mutants, the nuclear entry of PER-TIM heterodimer is not affected and proceeds as normal around midnight. Once in the nucleus, TIM(S1404A) interacts with the nuclear export machinery with higher affinity as compared to TIM(WT), leading to increased nuclear export and higher percentage of PER-TIM heterodimers in the cytoplasm. As a result, CLK phosphorylation in the nucleus is reduced as there are less PER-TIM heterodimers available to serve as scaffolds to recruit CLK kinases.

In agreement with this model, we observed dampening of daily CLK phosphorylation rhythms in *tim*(S1404A) flies (Fig. 5B). Specifically, there is significantly lower level of hyperphosphorylated CLK isoforms in late night to early morning. Given that CLK hyperphosphorylation is linked to reduced transcriptional activity, it is somewhat surprising that reduced CLK phosphorylation at night (ZT20 to ZT24) did not significantly enhance clock gene expression (Fig. 5A). This supports that other modifications, such as ubiquitination, are also important for regulating CLK transcriptional activity (71). USP8 has been shown to deubiquitylate CLK at late night to facilitate repression of clock genes.

We propose that reduced CLK hyperphosphorylation represents the key driver for the short period phenotype of *tim*(S1404A) mutant. Since CLK is not hyperphosphorylated by kinases recruited by the PER-TIM heterodimers at night, it does not have to be dephosphorylated by the CKA-PP2A complex (68) in the morning of the next cycle to activate clock gene transcription. This would explain the phase advance of CLK occupancy on circadian promoters (Fig. 5D) and CLK-activated *per* and *tim* expression (Fig. 5A). It is interesting to note that we did not observe diminished repression of clock genes despite the reduction of nuclear PER-TIM heterodimers at night (Fig. 5A), indicating additional mechanisms, e.g. chromatin remodeling and protein modifications, that are absent during this time of the circadian cycle are necessary to activate clock gene expression even when the level of PER repressor is reduced.

Curiously, *tim*(S1404D) mutants also exhibit a short period phenotype in behavioral rhythms (Fig. 1B). The fact that *tim*(S1404D) mutants display the opposite molecular phenotype in the context of nucleocytoplasmic localization of TIM and PER when compared to *tim*(S1404A) flies support that the S1404D mutation is phosphomimetic (Fig. 2 and 4). The short period phenotype of the *tim*(S1404D) mutant is therefore best explained by phase advance of the nuclear accumulation of PER-TIM heterodimer due to sustained inhibition of TIM-XPO1 interaction, which resulted in phase advance of subsequent events including CLK phosphorylation rhythm and PER and TIM turnover (Fig. 4F, 4G and S6).

Previous studies suggested that TIM phosphorylation plays a role in regulating its light-dependent degradation as well as its subcellular localization. Phosphorylation of tyrosine (pY) was first proposed to precede the degradation of TIM upon light exposure (40). We did not recover any pY residues in our MS analysis of PER-bound TIM. Nevertheless, we cannot rule out that Y phosphorylation may occur when TIM is not in complex with PER. Furthermore, it is possible that pY may result in the disassembly of the PER-TIM heterodimer or could result in very unstable TIM proteins that are difficult to purify from fly tissues. Instead of potential pY residues, we were able to identify a number of phosphorylated S/T residues that significantly impact behavioral phase shift responses to a light pulse at ZT15 or ZT21, suggesting they may be involved in mediating light-dependent TIM degradation (Fig. S1B and S1C). TIM(S568) is located within a functional NLS (23). Non-phosphorylatable *tim*(S568A) mutants exhibited arrhythmic locomotor activity rhythms (Fig. 1B), which is consistent with the phenotype of mutants defective in TIM nuclear entry (22, 23) and/or light entrainment (22). Future investigations are necessary to determine if S568 phosphorylation may regulate TIM subcellular localization and/or light responses. It will be also interesting to determine whether phosphorylation at S1389, S1393 and S1396, which are close to the NES identified in this study, play a role in regulating TIM nuclear export. Of note, we did not observe phosphorylation at T113, S297/T301 and T305/S309/S313 residues in our MS analysis. These residues were previously suggested to promote TIM nuclear entry when phosphorylated (22, 31). It is possible that these residues are more highly phosphorylated when TIM is not bound to PER.

In summary, we describe a phosphorylation-dependent nuclear export mechanism that regulates the nuclear accumulation of PER-TIM heterodimers and consequently the phase of CLK transcriptional activity in the molecular clock. We identified an NES motif in the previously characterized TIM CLD domain and showed that S1404 phosphorylation adjacent to this NES can regulate PER-TIM nuclear export. NES has been shown repeatedly to be an important regulatory motif that modulates localization and activities of transcriptional repressor in eukaryotes (72–76). The NES and S1404 module at the C-terminus of TIM is highly conserved in Drosophilids and in most species that have the *timeless* gene (Fig. S2A and S2B). Together with our MS analysis showing the phosphorylation of *Danaus plexippus* TIM(S1174), the homologous site of *Drosophila melanogaster* TIM(S1404) (Fig. S2C), we expect this phosphorylation-dependent mechanism that regulates TIM function to be conserved in insects. Interestingly in *timeout/timeless 2* (homolog of mammalian *timeless*), the ancestral paralog of *tim* that has a role in circadian photoreception but not in the oscillator itself (77, 78), this C-terminal NES is absent and serine is replaced by a glutamic acid (E) (Fig. S2B). We speculate that the gain of the NES and TIM(S1404) module at some point in evolution likely enabled TIM to cycle between subcellular compartments in a phosphorylation- and phase-dependent manner over the circadian cycle. This would allow CK2-dependent TIM phosphorylation to facilitate the phase-specific functions of PER-TIM heterodimers in specific lineages.

Finally, it is interesting to point out that a mutation in mammalian *timeless* that results in decreased nuclear TIM accumulation also leads to phase advance of human sleep-wake behavior, an output of the circadian clock (79). The circadian period length of mice expressing the TIM(R1081X) mutation, which manifests into human familial advanced sleep phase syndrome (FASPS), as determined by activity rhythm is identical to wild type mice. However, proliferating embryonic fibroblasts derived from heterozygous TIM(R1081X) mutant mice as well as mammalian U2OS and HEK293 cells expressing this same mutation exhibit a significant period-shortening. This suggests that decreased TIM nuclear accumulation in flies and mammals results in similar outcomes in the context of the molecular clock. Our results highlight an unexpected parallel between the functions of *Drosophila* and mammalian *timeless* in the molecular clock, even though the exact mechanisms and sequence motifs regulating their functions might have diverged. Analysis of *Drosophila tim* mutants could provide insights into the mechanisms that regulate the nuclear accumulation of mammalian TIM and further elucidate its functions in the mammalian clock (80). For instance, similar to what we deduced from *Drosophila tim*(S1404A) mutant, the advanced sleep phenotype in human FASPS R1081X patients could be the result of altered phosphorylation profile of BMAL1-CLOCK, leading to phase advance in circadian transcriptional activation.

## Materials and Methods

Extended materials and methods are provided in SI Materials and Methods.

## Data availability statement

All relevant data are within the manuscript and the SI. The *Drosophila* MS data has been deposited into Chorus repository (project ID 1424) and the *Danaus plexippus* DpNI MS data will be deposited into ProteomeXchange prior to publication.

## Acknowledgments

We thank Patrick Emery for providing *p{tim(WT)-luc}* transgene, α-PER and α-TIM antibodies for immunofluorescence, Steven Reppert and Christine Merlin for providing *Danaus plexippus* DpN1 cell line and pBA-*dpper*-FLAG plasmid, Carrie Partch for pHis::Parallel1 plasmid for protein expression. We thank Chris Fraser and Nancy Villa for their technical help on CLOCK antigen production. We thank the Bloomington *Drosophila* Stock Center and Vienna *Drosophila* Resource Center for providing fly stocks, and the Developmental Studies Hybridoma Bank for supplying α-PDF. The Confocal Microscopy facility was supported by NIH GM122968 to Pamela C. Ronald at UC Davis. Research in the laboratory of JCC is supported by NIH R01 GM102225, NIH R01 DK124068, and NSF IOS 1456297.

## Supplemental Information

## SI Materials and Methods

### Transgenic *Drosophila* construct design and fly transformation

A *p*{*tim*(WT)*-luc*} transgene, containing 4.1kb of the *tim* promoter, *tim* full-length coding region, and a luciferase reporter in the *pattB* vector, was kindly provided by Patrick Emery. The luciferase reporter was removed using MluI/XhoI restriction sites, and a 3XFLAG-6XHIS epitope was added in frame to the C-terminus of the *tim* coding region. To generate flies expressing non-phosphorylatable (Serine/Threonine (S/T) to Alanine (A)) or phosphomimetic (S/T to Aspartic acid (D)) *tim* mutants, pAc-*ls-tim*-HA was used as the template for site-directed mutagenesis using Pfu Turbo Cx DNA polymerase (Agilent Technologies, Santa Clara, CA) (see Table S3 for mutagenic primer sequences). After mutagenesis and confirmation by Sanger sequencing (UC Davis DNA Sequencing Facility), the mutant variants of 2.8 kb MluI-XbaI *tim* subfragments were used to replace the corresponding WT fragments in *pattB*-*p*{*tim*(WT)*-*3XFLAG-6XHIS}. PhiC31 site-directed recombination was used for transgenesis (1). Plasmids were injected into *yw* fly embryos carrying *attP* sites on chromosome 3 (*attP2*) (BestGene, Chino Hills, CA). Transformants were crossed with *yw*; *tim^0^* flies (2) to remove endogenous copies of *tim* prior to behavioral and molecular analyses.

Targeted expression of *ck2α* dsRNA in *tim*-expressing neurons was achieved via the UAS/Gal4 system (3). The Gal4 driver line, *w; UAS-dicer2; tim-UAS-Gal4* (*TUG*) (4), was used to drive expression in *tim* expressing clock neurons. *UAS-ck2α^tik^* responder line (B24624) from the Bloomington *Drosophila* Stock Center was used to reduce endogenous function of CK2α.

### Identification of TIM phosphorylation and O-GlcNAcylation sites from fly tissues

PER-bound TIM phosphorylation and O-GlcNAcylation sites were identified from the label-free mass spectrometry (MS) proteomics experiments as previously described (5). Procedures for immunoprecipitation of PER-TIM complexes and mass spectrometry (MS) were previously described (5). Epitope-tagged PER proteins were pulled down using α-FLAG and PER-bound TIM proteins were pulled down simultaneously and subjected to MS analysis.

Mass spectrometric data was analyzed with PEAKS Studio X+ (Bioinformatics Solutions Inc., Canada). Raw data refinement was performed with the following settings: Merge Options: no merge, Precursor Options: corrected, Charge Options: 1-6, Filter Options: no filter, Process: true, Default: true, Associate Chimera: yes. *De novo* sequencing and database searching were performed with a Parent Mass Error Tolerance of 10 ppm. Fragment Mass Error Tolerance was set to 0.02 Da, and Enzyme was set to none. The following variable modifications were applied: Oxidation (M), pyro-Glu from Q (N-term Q), phosphorylation (STY), acetylation (protein N-terminal) and HexNAc (STNY). Carbamidomethylation (C) was set as fixed modification. A maximum of 5 variable PTMs were allowed per peptide. A custom database of appropriate size (550 protein sequences) containing TIMELESS protein sequence (UniProt ID A0A1W5PW00) from UniProt was used for database searching. Database search results were filtered to 1% PSM-FDR. Phosphosite localization was validated by inspecting fragment ion spectra of all phosphopeptides. Identification of GlcNAc-modified peptides was confirmed by inspecting the corresponding fragment ion spectra for the presence of characteristic fragment ions (m/z 168.07, 186.08, 204.09). The *Drosophila* MS data has been deposited into Chorus repository (project ID 1424).

### *Danaus plexippus* DpN1 cell culture

Monarch butterfly DpN1 cells (6), kindly provided by Steven Reppert and Christine Merlin, were grown at 28°C in Grace’s medium (Thermo Fisher Scientific, Waltham, MA), supplemented with 10% Fetal Bovine Serum (FBS) (VWR, Radnor, PA) and 1X penicillin/streptomycin (Thermo Fisher Scientific). Cells were passed every 7 days. Old medium was removed and cells were washed with cell culture grade 1XPBS (Thermo Fisher) once before treating with Trypsin/EDTA (Thermo Fisher) for at least two minutes at 28°C. To halt trypsinization, FBS-containing Grace’s medium was added to the cells. Cell suspensions were then passed into new tissue culture flask with FBS-containing Grace’s medium.

### Affinity purification of *Danaus plexippus* PER (dpPER) followed by mass spectrometry

Affinity purification was performed as previously described (5) with the following modification. For each time-point, roughly 2.5g of cell pellet in Lysis Buffer (20mM Hepes pH 7.5, 1mM DTT, 0,5mM PMSF and SIGMAFAST EDTA-free protease inhibitor cocktail were dounced prior to centrifugation at 800xg for 15 minutes at 4°C to separate nuclear and cytoplasmic fractions. The nuclear fraction (pellet) was washed twice with Lysis Buffer prior to resuspension in Nuclear Extraction Buffer (20mM Tris-HCl pH 7.5, 150mM NaCl, 0,5mM EDTA, 1mM DTT, 1mM MgCl_2_, 1% Triton X-100, 0.4% sodium deoxycholate, 0.5mM PMSF, 10mM NaF, 10% glycerol and SIGMAFAST) and dounced with tight pestle supplemented with MG132 (Sigma) and DNase (Promega, Madison, WI). The cytoplasmic fraction (supernatant) was supplemented with 150mM NaCl, 1mM MgCl_2_, 0.5mM EDTA, 1% Triton X-100, 0.4% sodium deoxycholate, 10mM NaF, 10% glycerol and SIGMAFAST. After 30 minutes incubation at 4°C, nuclear fraction was diluted to 0.1% SDS with concentration of other content unchanged. Nuclear and cytoplasmic fractions were then centrifuged at 15,000 rpm for 15 minutes. Supernatants were incubated with gammabind sepharose beads (GE Healthcare, Piscataway, NJ) for 30 minutes at 4°C to reduce nonspecific binding prior to overnight incubation with α-dpPER (GP5913, RRID: AB_2832970). On the second day, samples were incubated with gammabind sepharose beads (GE Healthcare). Beads were washed three times with Wash buffer (20mM HEPES, 150mM NaCl, 1mM MgCl_2_, 1mM DTT0.5mM EDTA, 1% Triton X-100, 0.4% sodium deoxycholate, 10mM NaF, 0.5mM PMSF, 10% glycerol) and subsequently eluted in 200ul 2X SDS sample buffer at 95°C for 4 minutes. After resolving eluates on a Tris-Tricine gel, excised gel containing eluates was digested with protease and followed by mass spectrometry as previously described (5). DpnI1 MS data will be deposited into ProteomeXchange prior to publication.

### Locomotor activity assay

Daily locomotor activity rhythms in male flies was assayed using the *Drosophila* Activity Monitoring System (DAMS, Trikinetics, Waltham, MA) as described previously (7). Flies were entrained for 4 days in light/dark (LD) cycles (12h light/12h dark), followed by 7 days of constant darkness (DD) to assess their free-running rhythms at 25 ̊C.

### Assaying responses to light pulse

Male flies were entrained for 4 days in LD cycle (12h light/12h dark). In the dark phase on LD4, the light-pulsed (LP) flies were given a 10-minute pulse of light at ZT 15 or ZT 21 before being placed in 7 days of DD, while the non-light pulsed (NLP) flies were not exposed to light pulse treatments. Activity rhythms were measured using the DAMS and analyzed as using FaasX as previously described (7). Peaks in activity rhythms were restricted to between ZT 6 and ZT 18 and converted into a value in degrees using Excel (Microsoft, Redmond, WA) (24 hours = 360°). LD3 data was used to normalize the DD1 data of each fly by subtracting the LD3 from DD1, within each respective genotype. *tim*(WT) degree values were then subtracted from each mutant to determine the phase shift as a result of the light pulse. The difference in degrees was then converted back to hours using Excel (Microsoft).

### Plasmids for *Drosophila* S2 (Schneider 2) cell culture

pAc-*per*-V5-HIS and pAc-*tim*-HA were described previously (8). *Exportin 1* (*xpo1*) ORF was amplified from cDNA that was reverse transcribed (Superscript IV, Thermo Fisher Scientific) from total RNA extracted from fly heads using TRI Reagent (Sigma, St. Louis, MO). The PCR product was cloned into pAc-3XFLAG-6XHIS (9).

### *Drosophila* S2 cell culture and transfection

*Drosophila* S2 cells and *Schneider’s Drosophila* medium were obtained from Life Technologies (Carlsbad, CA). For all cell culture experiments, S2 cells were seeded at 1 × 10^6^ cells/ml in a 6-well plate and transfected using Effectene (Qiagen, Germantown, MD). For coimmunoprecipitation (coIP) assays, S2 cells were co-transfected with 0.8μg of pAc-*per*-V5-HIS (herein referred to as pAc-*per*-V5), 0.8μg of pAc-*xpo1*-3XFLAG-6XHIS (herein referred to as pAc-*xpo1*-FH) and 0.8μg of pAc-*tim*(X)-HA, where X is either WT or S1404A to detect protein-protein interactions. In control IPs to detect non-specific binding, cells were co-transfected with pAc-*per*-V5-His and pAc-*xpo1*-FH without pAc-*tim*(X)-HA. S2 cells were harvested 40 hours after transfection. For IP to detect TIM(pS1404), S2 cells were co-transfected with 0.8μg of pAc-*tim*(X)-HA and either 0.2μg of pMT-*ck2α*-V5 or 0.2μg pMT-V5-His as empty control. Expression of *ck2α* was induced with 500 μM CuSO4 immediately after transfection and cells were harvested 40 hours after kinase induction.

### Coimmunoprecipitation (CoIP) in *Drosophila* S2 cells

CoIP experiments were performed as described previously (10) with the following modifications. Cells were harvested 40 hours after transfection and lysed with modified RIPA (20mM Tris-HCl pH 7.5, 150mM NaCl, 10% glycerol, 1% Triton X-100, 0.4% sodium deoxycholate, 0.1% SDS) supplemented with 1mM EDTA, 25mM NaF, 0.5mM PMSF, and SIGMAFAST. Proteins were incubated with 20μl α-HA or α-FLAG M2 resins (Sigma) to pull down TIM or XPO1, respectively. Immune complexes were analyzed by Western blotting. Signal intensity of interacting protein was normalized to the intensity of the bait protein.

### Western blotting and antibodies

Protein extractions from *Drosophila* S2 cells and adult fly heads, western blotting, and image analysis were performed as previously described (10, 11). Primary antibodies: α-HA 3F10 (Roche, Indianapolis, IN) at 1:2000 for TIM-HA, α-V5 (Thermo Fisher Scientific) at 1:3000 for PER-V5, α-FLAG (Sigma) at 1:7000 for XPO1-FLAG, α-TIM (R5839, RRID:AB_2782953) at 1:1000 for TIM (12), α-pS1404 (RB S4602-2, RRID:AB_2814716) at 1:2000 for TIM(pS1404) isoforms, α-CLK (GP6139, RRID:AB_2827523) at 1:2000 for CLK, α-PER (GP5620; RRID:AB_2747405) at 1:2000 for PER and α-HSP70 (Sigma) at 1:10000 was used for to indicate equal loading and for normalization. Secondary antibodies conjugated with HRP were added as follows: α-mouse IgG (Sigma) at 1:2000 for α-V5 detection, 1:2000 for α-FLAG detection, or 1:10,000 for α-HSP70 detection, α-guinea pig IgG (Sigma) at 1:1000 for α-PER detection, α-rabbit IgG (Sigma) at 1:2000 for α-pS1404 detection ,and α-rat IgG (Sigma) at 1:1000 for detecting α-HA and α-TIM.

### Generating *Drosophila* CLOCK antibodies

The first 1770 nucleotides of *Drosophila clk* cDNA (Flybase: FBpp0099478) was subcloned into a modified His-tagged pFastBac1 vector (Invitrogen, Carlsbad, CA) as previously described (13). The recombined construct, pFastBac1-6XHis-d*clk* (1-1770), was transformed into DH10BAC *E. coli* (Invitrogen) and bacmid DNA was then purified. To generate viral stock, bacmid DNA was transfected into Sf9 cells using XtremeGENE 9 transfection reagent (Sigma) and media is collected according to the manufacturer’s protocol. Viral stock was used to infect High Five cells (Thermo Fisher Scientific) for large-scale expression of CLK antigen. As the CLK antigen is insoluble in extraction buffer (20mM Hepes pH 7.5, 400mM KCl, 5mM imidazole, 10% glycerol, 10mM β-mercaptoethanol) supplemented with 1X SIGMAFAST (Sigma), we collected the insoluble cell pellet after extraction and dissolved it in denaturing solution (50mM Na3PO4, 1% SDS) by boiling for 10 minutes. SDS in the sample was diluted to 0.05% before purification using 5ml of Ni-NTA Superflow nickle-charged resin (Qiagen). CLK antigen was sent to Covance Inc. (Princeton, NJ) for antibody production in guinea pigs.

### Generating *Danaus plexippus* PERIOD antibodies

*Danaus plexippus per* cDNA sequence that encodes amino acid 898-1095 was amplified from pBA-*dpper*-FLAG (6) and subcloned into the pHis::Parallel1 bacterial expression vector kindly provided by Carrie Partch. This plasmid was transformed into BL21 (DE3) *E. coli* strain and protein expression was induced as described previously (14). dpPER fragment was affinity-purified with a 5ml IMAC Nickle column using the NGC system (Bio-Rad, Hercules, CA) and used as immunogen in guinea pigs (Covance).

### Generation of TIM(S1404) phosphospecific antibodies

Phosphospecific antibodies were generated by ProteinTech Group, Inc (Rosemont, IL). Rabbits were immunized with a 15-amino-acid peptide (amino acid 1397-DLTRMYVpSDEDDRLE-amino acid 1411; where pS= phosphoserine). Immunoblotting analysis was performed as previously described (11) with modifications detailed in the “Western blotting and antibodies” section.

### Immunoprecipitation (IP) and phosphatase (λ*-PP*) treatment to detect phosphorylated TIM(S1404) in *Drosophila* S2 cells and transgenic flies

IP and λ-PP treatment were performed as described previously (11). TIM proteins from S2 cell extracts were pulled down using 20μl of α-HA resin per IP reaction. IP with fly extracts were performed as previously described (11) using 4μl α-TIM and 20ul gammabind sepharose beads (GE Healthcare) per IP reaction.

### Immunofluorescence and confocal imaging of *Drosophila* clock neurons

Brain dissections and immunofluorescence staining procedures were performed as described previously (15). Briefly, 3-5d old flies were entrained for 4 days and fixed with 4% paraformaldehyde at 2-hr intervals between ZT16 and ZT22. Brains were washed three times in 1XPBST (0.1% Triton X-100 in PBS), blocked with 10% Normal Goat Serum (Jackson Immunoresearch, West Grove, PA) in PBST and incubated with primary antibodies overnight. Primary antibodies against PDF, PER, and TIM were used at the following dilutions: 1:1500 rabbit α-PER (Gift from Dr. Patrick Emery) (16), 1:100 GP α-TIM (Gift from Dr. Patrick Emery) (17) and 1:400 mouse α-PDF (C7-C; Developmental Studies Hybridoma Bank, Iowa City, IA). Brains were then washed and probed with secondary antibodies at a 1:200 dilution for α-rat-cy3 (Jackson Immunoresearch, 706-165-148), α-rabbit IgG Alexa Fluor 488 (Jackson Immunoresearch, 711-545-152) and α-mouse-cy5 (Jackson Immunoresearch, 715-175-150). Eight to ten fly brains for each genotype were dissected and imaged. Representative images are shown. Fiji software was used for image analysis (18).

### Quantitative RT-PCR

Extraction of total RNA from fly heads, reverse transcription, and quantitative real time PCR were performed as previously described (10). Primers for qPCR were also described previously (10).

### Chromatin Immunoprecipitation (ChIP)

CLK-ChIP was performed as described previously (10). Primers for *per* CRS and *tim* E-box were described previously (10). The average of ChIP signals for two intergenic regions, one on chromosome 2R (Table S3) and one on the X chromosome (10), was used for non-specific background deduction.

### Statistics

RAIN, DODR, Rayleigh test, Shapiro-Wilk normality test and Watson Williams test were performed in R (19–22). Other statistical analyses were performed using GraphPad Prism 8.0 (GraphPad Software, La Jolla, California). In the case of normally distributed data (Shapiro-Wilk normality test, p>0.05), two-way ANOVA was performed if more than two groups were compared; two-tailed Student’s t test were performed if only two groups were compared. If data were not normally distributed, non-parametric Kruskall-Wallis test was applied. Asterisks indicate significant differences in mean values between genotypes or conditions at indicated time-points.

## SI Figure Legends

**Figure S1.**
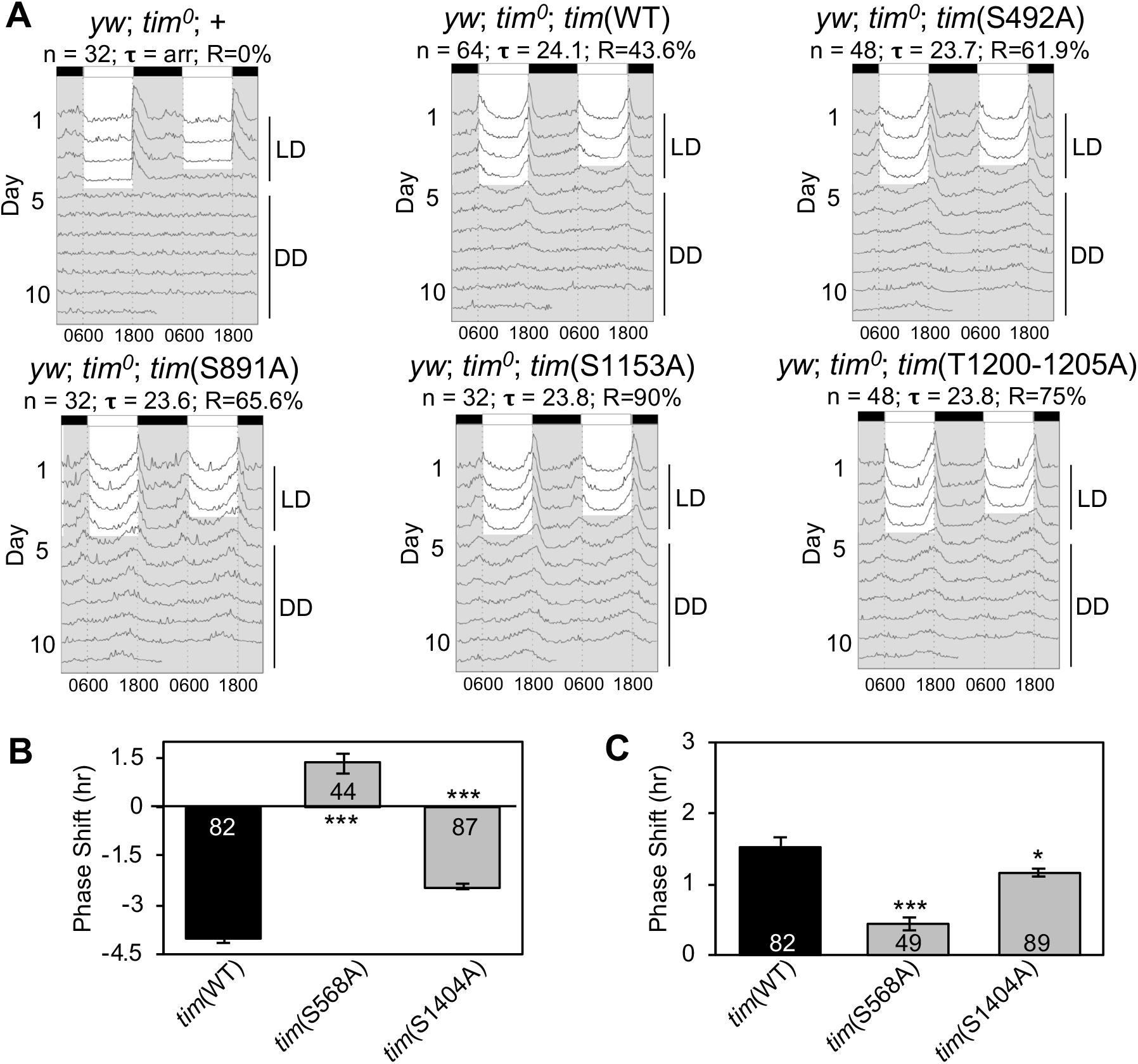
Daily locomotor activity rhythms and responses to light pulse are altered in TIM phosphorylation site mutants. (A) Double-plotted actograms of *yw*; *tim^0^* flies carrying transgenes for site-specific TIM phosphorylation mutation. n represents the sample size for behavioral assay. Tau (**τ**) represents the average period length of the group of flies in DD. R represents percentage of flies that are rhythmic. Flies were entrained for 4 days in LD cycles and then switched to 7 days of constant darkness, DD. (B-C) Bar graphs showing the phase shift of *tim*(WT) and *tim* mutants in response to light pulse at (B) ZT15 and (C) ZT21, respectively. Error bars indicate SEM; ***p<0.001, *p<0.05, as compared to *tim*(WT), two-tailed Student’s t test. Sample size (n) is shown at the bottom of each bar. The Rayleigh test was used to confirm significant synchronization of *tim*(WT) and *tim*(S1404A) fly populations on LD3 and DD1 (p<0.0001). Behavioral arrhythmicity of *tim*(S568A) was confirmed by Rayleigh test (p=0.6105). The Watson-Williams test was used to confirm a significant phase shift (p<0.01) for *tim*(S1404A) mutants.

**Figure S2.**
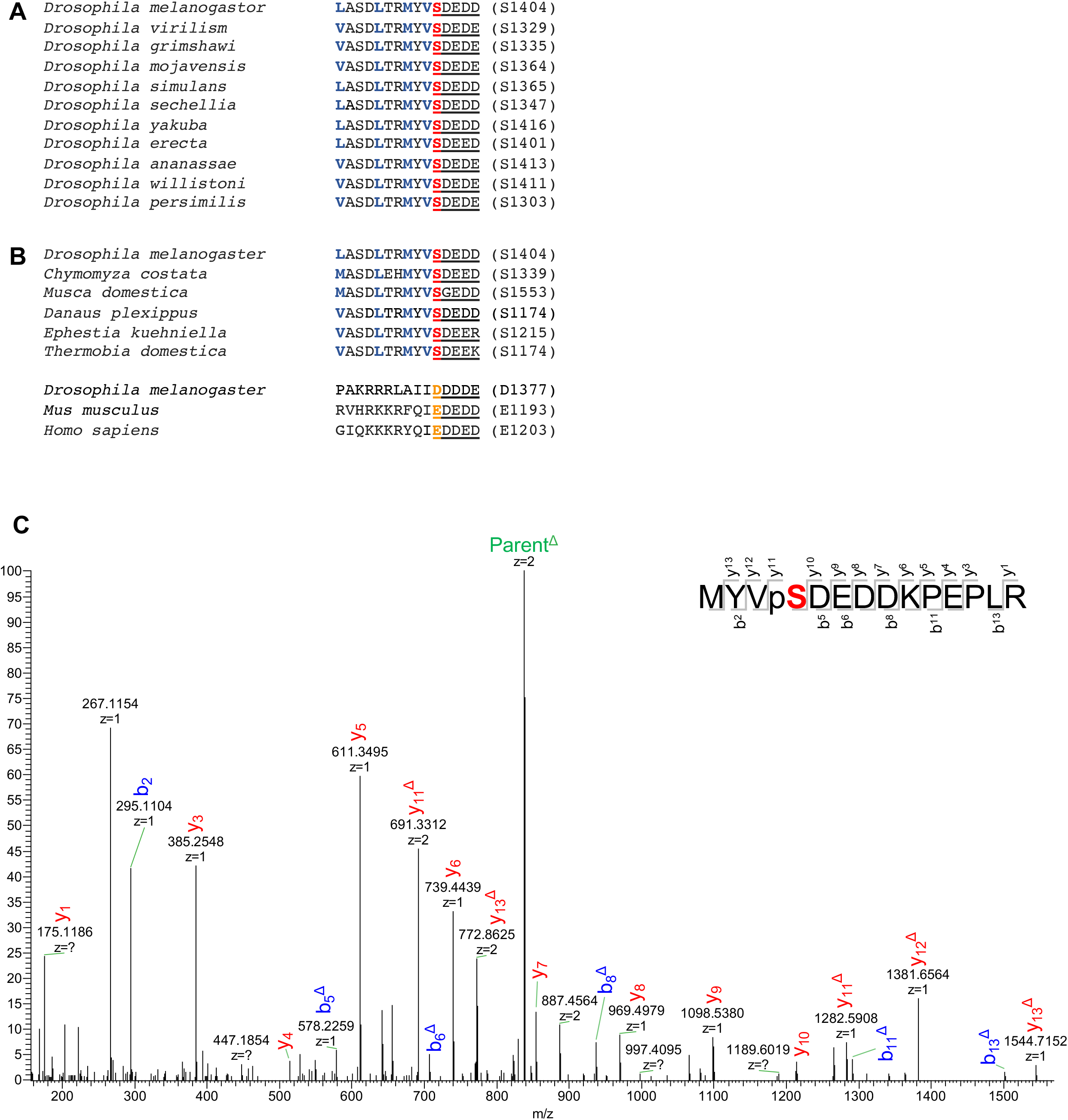
TIM(S1404) phosphorylation is conserved in insects. (A) Amino acid sequences surrounding TIM(S1404) are conserved in *Drosophila* species. *D. pseudoobscura* is not included because it is missing the S1404 region. S1404 is shown in red. Critical hydrophobic amino acids within NES sequence motif are shown in blue. Consensus CK2 site is underlined. (B) Alignment of *D. melanogaster* TIM(S1404) region to the corresponding homologous sequence in indicated insect species, mouse (*Mus musculus*), and human (*Homo sapiens*). Glutamic acid or aspartic acids in *D. melanogaster* TIMEOUT/TIMELESS2, mouse and human TIMELESS sequences that potentially replaces TIM(S1404) in *D. melanogaster* TIMELESS are in orange. (C) Phosphorylation at S1404 homologous residue in *Danaus plexippus*, dpTIM(S1174), is detected at ZT16, ZT20 and ZT24 in DpN1 cells by mass spectrometry proteomics. Representative HPLC/MS/MS spectrum showing phosphopeptide 1171-MYVpSDEDDKPEPLR-1184, where pS= phosphoserine. Parent^Δ^ denotes neutral loss ion. The notations bnΔ or ynΔ denotes the corresponding bn or yn ions with neutral loss.

**Figure S3.**
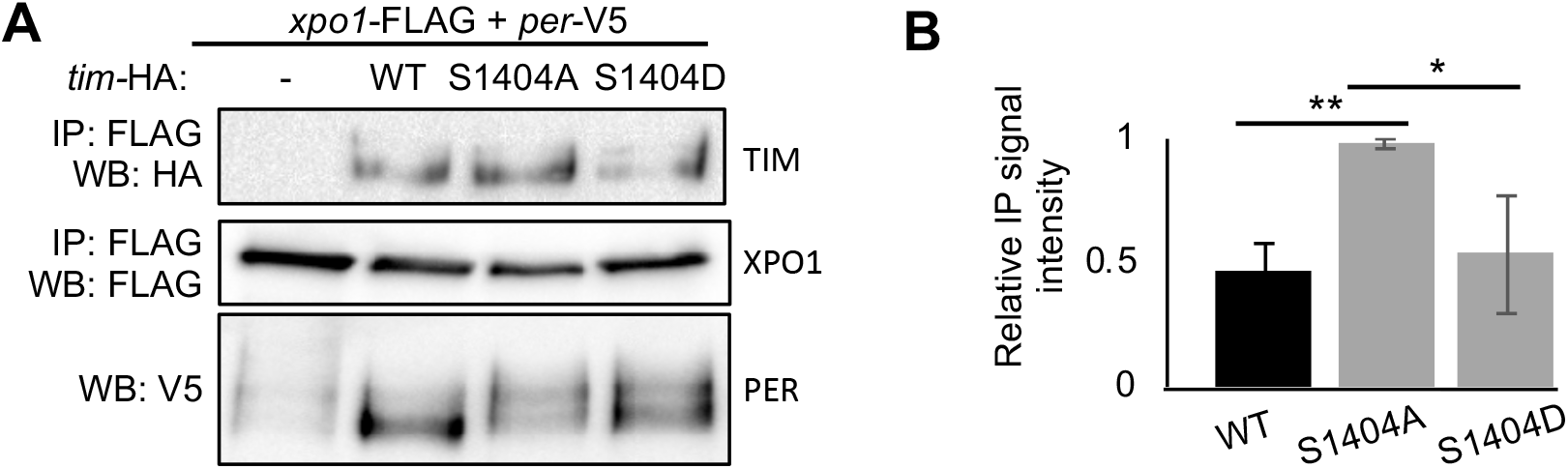
TIM(S1404D) compromises TIM-XPO1 interaction. (A) Western blots showing coimmunoprecipitation (coIP) to examine the interactions of TIM(WT), TIM(S1404A) or TIM(S1404D) to XPO1 in *Drosophila* S2 cells expressing pAc-*xpo1*-3XFLAG-6XHIS and pAc-*per*-V5 in the presence or absence of pAc-HA plasmids expressing *tim* variants. Protein extracts were directly analyzed by immunoblotting (α-V5 for PER) or immunoprecipitated with α-FLAG resins and analyzed by immunoblotting to detect baits and interactors. (B) Bar graph displaying quantification of (A). Values for binding are normalized to amount of bait detected in the IPs and expressed as relative signal intensity (high value = 1). Error bars indicate ± SEM (n=3), **p<0.01, *p<0.05, one-way ANOVA.

**Figure S4.**
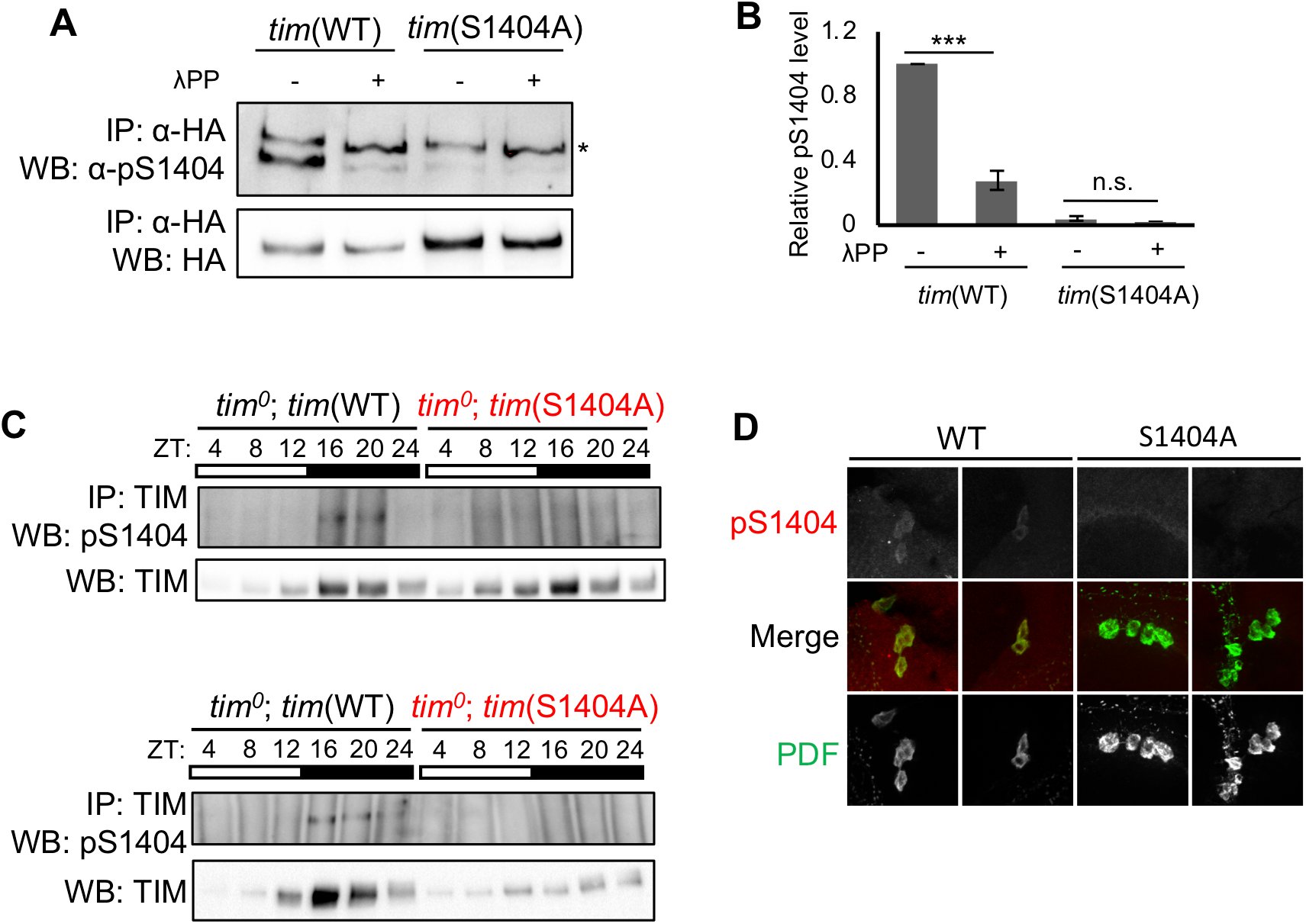
TIM(S1404) is phosphorylated in wild type flies between mid to late night. (A) *Drosophila* S2 cells were transfected with pAc-*tim*(WT)*-*HA or pAc-*tim*(S1404A)*-*HA. Protein extracts were incubated with α-HA resins. Half of the immunocomplexes received lambda phosphatase (λPP) treatment while the other received sham treatment. TIM(pS1404) and TIM protein levels were analyzed by Western Blotting. Asterisk (*) indicates nonspecific signal (upper band). (B) Quantification of TIM(pS1404) was normalized to total TIM isoforms. Error bars indicate ± SEM (n=2), ***p<0.001, one-way ANOVA. (C) Fly heads of the specified genotypes were collected at the indicated times on LD3 after 2 days of LD entrainment. TIM was immunoprecipitated with α-TIM prior to western blotting with α-pS1404 (top panel). Total TIM isoforms are shown in the bottom panel. 2 biological replicates are shown. (D) Representative confocal images of LNvs clock neurons in adult fly brains stained with α-TIM(pS1404) (red) and α-PDF (green). Single channels are shown in grey scale. Flies were entrained for 2 days in LD cycles and collected at ZT16 on LD3, fixed and analyzed by immunofluorescence and confocal imaging.

**Figure S5.**
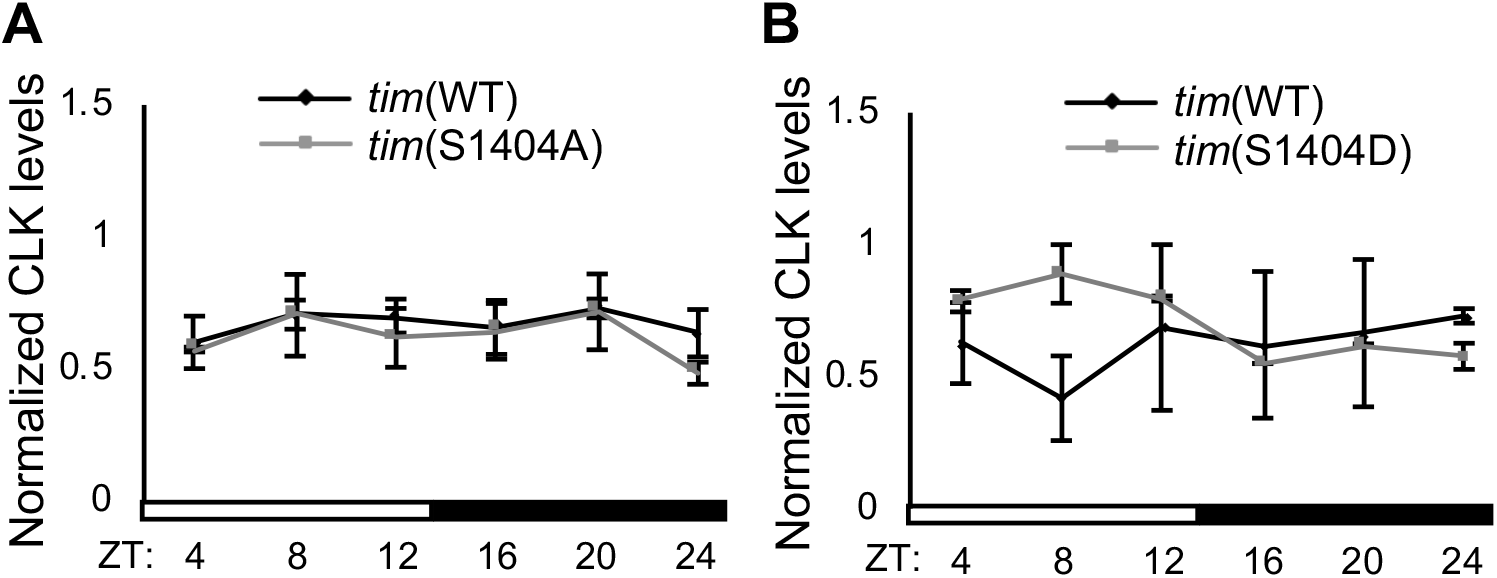
*tim*(S1404) mutations do not alter daily rhythms in CLK protein abundance in fly heads. (A) Quantification of CLK abundance presented in Figure 5B; (B) Quantification of CLK in Figure S6B. Error bars indicate ± SEM (n=3), p>0.05 at all ZTs, two-way ANOVA.

**Figure S6.**
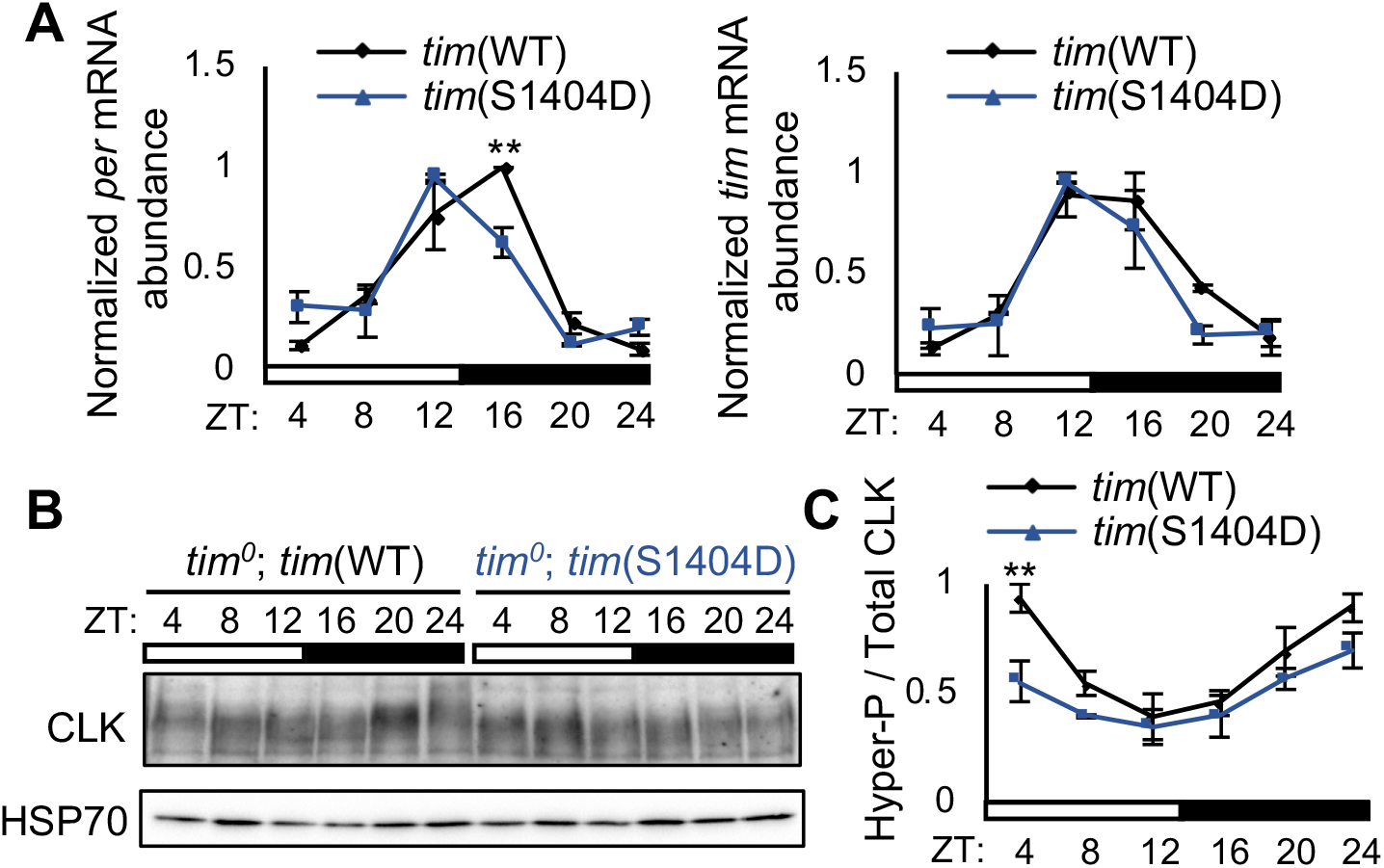
Increased TIM nuclear retention in *tim*(S1404D) mutant leads to shortened molecular rhythms. (A) Steady state mRNA expression of *per* and *tim* in heads of *tim*(WT) and *tim*(S1404D) flies, entrained in LD cycles and assayed on LD3 (n=3). (B) Western blots comparing CLK profiles in heads of *tim*(WT) and *tim*(S1404D) entrained and collected as in (A). ⍺-HSP70 was used to indicate equal loading and for normalization. (C) Quantification of hyperphosphorylated/total CLK ratios. The top half of the CLK signal shown at ZT24 in *tim*(WT) flies (lane 6) was used as a reference to denote hyperphosphorylated CLK isoforms. Error bars indicate ± SEM (n=3). **p<0.01, two-way ANOVA.

---

**Table S1.**
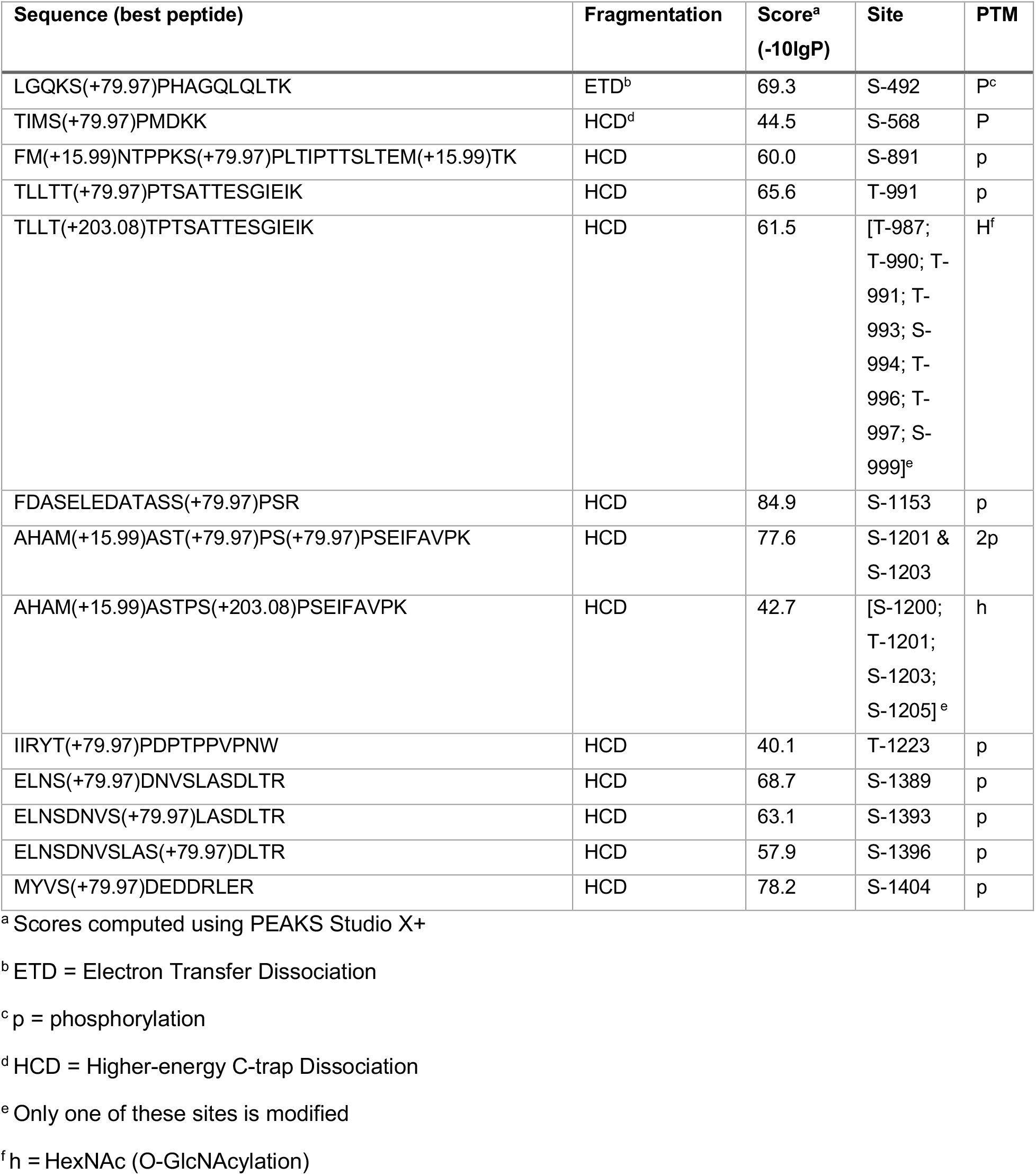
Identification of TIM phosphorylation and O-GlcNAcylation sites in *Drosophila* head tissues.

**Table S2.** Daily locomotor activity rhythms of *tim* mutants.

| Genotype | Period (h)<br>(mean ± SEM) | Power<sup>a</sup> | Rhythmicity<br>(%)<sup>b</sup> | No. of<br>flies<br>tested | No of flies<br>surviving<sup>c</sup> |
| --- | --- | --- | --- | --- | --- |
| *yw; tim<sup>0</sup>; +* | AR<sup>d</sup> | ND<sup>e</sup> | 0 | 32 | 31 |
| *yw; tim<sup>0</sup>; tim(WT)* | 24.1 ± 0.12 | 56.4 | 43.6 | 64 | 55 |
| *yw; tim<sup>0</sup>; tim(S492A)* | 23.7 ± 0.06 | 88.6 | 61.9 | 48 | 42 |
| *yw; tim<sup>0</sup>; tim(S568A)* | 24.9 ± 0.43 | 63.3 | 15.4 | 32 | 26 |
| *yw; tim<sup>0</sup>; tim(S891A)* | 23.6 ± 0.08 | 88.2 | 65.6 | 32 | 32 |
| *yw; tim<sup>0</sup>; tim(S1153A)* | 23.8 ± 0.06 | 99.9 | 90 | 32 | 30 |
| *yw; tim<sup>0</sup>; tim(T1200-1205A)* | 23.8 ± 0.06 | 89.9 | 75 | 48 | 36 |
| *yw; tim<sup>0</sup>; tim(S1404A)* | 22.4 ± 0.06 | 83.2 | 68.3 | 64 | 60 |
| *yw; tim<sup>0</sup>; tim(S1404D)* | 22.4 ± 0.11 | 47.5 | 48.4 | 32 | 31 |

<sup>a</sup> Measures the strength or amplitude of the locomotor activity rhythm (in arbitrary units)
<sup>b</sup> Percentage of flies that are rhythmic
<sup>c</sup> Number of flies that survived until the end of the experiment
<sup>d</sup> AR denotes Arrhythmic
<sup>e</sup> ND denotes Not Determined

**Table S3:**
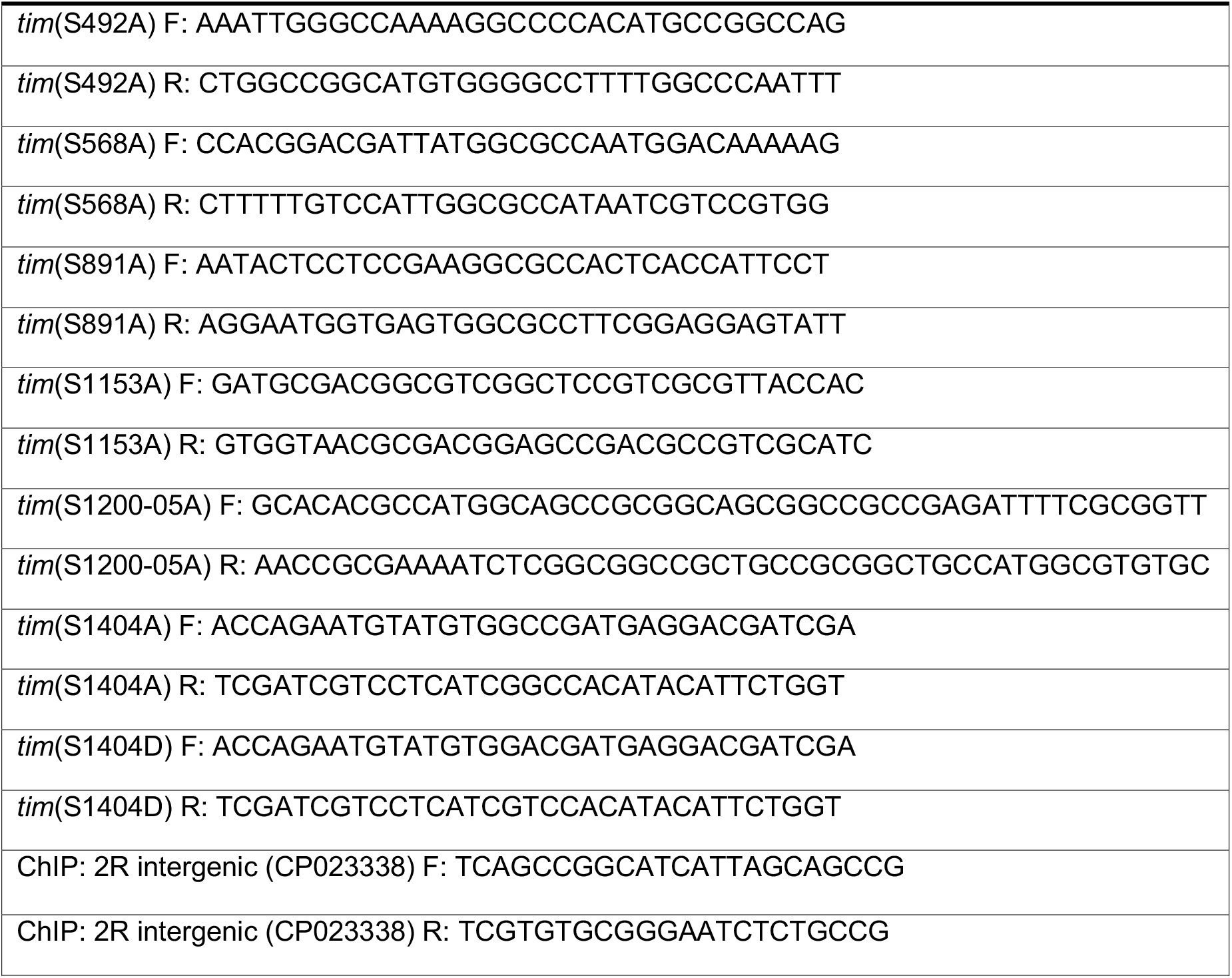
Primers for mutagenesis and ChIP.

